# KLF17 promotes human naïve pluripotency but is not required for its establishment

**DOI:** 10.1101/2020.12.18.423466

**Authors:** Rebecca A. Lea, Afshan McCarthy, Stefan Boeing, Kathy K. Niakan

## Abstract

Current knowledge of the transcriptional regulation of human pluripotency is incomplete, with lack of inter-species conservation observed. Single-cell transcriptomics of human embryos previously enabled us to identify transcription factors, including the zinc-finger protein KLF17, that are enriched in the human epiblast and naïve hESCs. Here we show that KLF17 is expressed coincident with the known pluripotency factors NANOG and SOX2 across human blastocyst development. We investigate the function of KLF17 in pluripotency using primed and naïve hESCs for gain- and loss-of-function analyses. We find that ectopic expression of KLF17 in primed hESCs is sufficient to induce a naïve-like transcriptome and that KLF17 can drive transgene-mediated resetting to naïve pluripotency. This implies a role for KLF17 in establishing naïve pluripotency. However, CRISPR-Cas9-mediated knockout studies reveal that KLF17 is not required for naïve pluripotency acquisition *in vitro*. Transcriptome analysis of naïve hESCs identifies subtle effects on metabolism and signalling following KLF17 loss of function, and possible redundancy with the related factor, KLF5. Overall, we show that KLF17 is sufficient, but not necessary, for naïve pluripotency under the given *in vitro* conditions.

**Summary statement:** Investigating KLF17 in human pluripotency reveals that it is sufficient, but not necessary, to establish naïve hESCs. We posit that KLF17 is a peripheral regulator, like KLF2 in the mouse.

## Introduction

Model organisms such as the mouse have allowed for the identification of molecular mechanisms that regulate early mammalian development (Rossant, 2016), some of which are conserved in humans (Gerri et al., 2020). Despite the continued importance of comparative studies in mouse and other organisms, some aspects of early development such as developmental timing, chromatin accessibility and transcription factor function are distinct compared to humans (Niakan and Eggan, 2013, Fogarty et al., 2017, Gao et al., 2018). In particular, the advent of single-cell sequencing technologies has allowed in-depth transcriptomic analysis of human embryos, revealing a number of molecular differences compared to the mouse (Yan et al., 2013, Blakeley et al., 2015, Petropoulos et al., 2016, Stirparo et al., 2018). Our previous analysis highlighted that a number of genes thought of as canonical pluripotency-associated factors in the mouse, including *KLF2, ESRRB* and *BMP4* (Blakeley et al., 2015), are not expressed in the pluripotent epiblast (EPI) of the human pre-implantation embryo, which forms the embryo proper. Conversely, we also highlighted a number of genes that are specifically enriched in the human EPI, but not expressed in the pluripotent cells of the mouse embryo, including transcriptional regulators and signalling components (Blakeley et al., 2015).

Of these human EPI-enriched genes, the zinc finger DNA-binding protein KLF17 has drawn considerable interest. KLF17 is one of 11 human paralogues of the Krüppel-like transcription factor family involved in development, which includes KLF4, a commonly used reprogramming factor (Takahashi and Yamanaka, 2006) and KLF2, a known pluripotency regulator in the mouse (Hall et al., 2009). Given the lack of *KLF2* expression in the human EPI, it is interesting to speculate that KLF17 might function in a similar way. Indeed, the expression patterns of *KLF2* and *KLF17* in the human embryo are diametrically opposite to those of *Klf2* and *Klf17* in the mouse embryo (Yan et al., 2013, Blakeley et al., 2015). While *Klf17* appears to be maternally deposited in the mouse zygote and its expression is abolished around the 8-cell stage, *KLF17* is dramatically upregulated in the 8-cell human embryo, following embryonic genome activation (EGA). Conversely, *Klf2* is expressed from the 2-cell stage, corresponding to mouse EGA, and continues through to the blastocyst stage, but human *KLF2* is expressed only pre-EGA, in the zygote to 4-cell embryo. The human KLF17 and KLF2 sequences share ~60% homology across the C-terminal region containing the functional C_2_H_2_-type zinc-finger domains. KLF17 and mouse KLF2 also have additional homologous regions (~50%) throughout the protein, including part of a region in mouse KLF2 annotated as a protein-protein interaction domain, which may contribute to regulation and/or functional specificity. Furthermore, in mouse embryonic stem cells (mESCs), the triple knockout of *Klf2, Klf4* and *Klf5* can be rescued by ectopic expression of human *KLF17*, but not mouse *Klf17* (Yamane et al., 2018). Finally, the human and mouse KLF17 protein sequences have less similarity overall than other pairs of KLF orthologues (van Vliet et al., 2006). This is all suggestive of rapid, divergent evolution of the human and mouse KLF genes and a potential switching of their function between species.

To date, KLF17 has primarily been studied in the context of cancer, where it has been implicated as a tumour suppressor by interacting with TGFβ/SMAD signalling (Ali et al., 2015b) and p53 (Ali et al., 2015a) and inhibiting epithelial-to-mesenchymal transition (Gumireddy et al., 2009, Zhou et al., 2016). Since the recognition of its human EPI-specific expression, KLF17 has been widely used as a marker of pluripotency in the human embryo (Blakeley et al., 2015, Guo et al., 2016, Shahbazi et al., 2017, Kilens et al., 2018). The expression of *KLF17* throughout pre-implantation development, and in particular in pluripotent cells, is also conserved in a number of other organisms, including non-human primates (rhesus monkey, *Macaca mulatta* (Wang et al., 2017); common marmoset, *Callithrix jacchus* (Boroviak et al., 2015); and cynomolgus monkey, *Macaca fascicularis* (Nakamura et al., 2016)), and pig (*Sus scrofa* (Bernardo et al., 2018, Ramos-Ibeas et al., 2019)). Intriguingly, KLF17 expression is not detectable in conventionally derived “primed” human embryonic stem cells (hESCs) (Blakeley et al., 2015, Stirparo et al., 2018), indicating that expression from EPI cells is lost during the derivation process. However, newer methods for deriving and/or culturing hESCs and human induced pluripotent stem cells (hiPSCs) in a naïve pluripotent state result in the maintenance or reinstatement of *KLF17* gene activity (Theunissen et al., 2014, Guo et al., 2017, Guo et al., 2016, Liu et al., 2017, Kilens et al., 2018). This pattern of expression suggests the intriguing possibility that KLF17 acts as a transcriptional regulator of human naïve pluripotency, as exhibited in the *bona fide* state of the pre-implantation EPI and approximated in the *in vitro* naïve hESC models. This hypothesis has also been explored by independent transcriptome analysis (Stirparo et al., 2018).

Studies to date have only conclusively shown that KLF17 is a marker of human pluripotency. Here, we set out to determine the function of KLF17, finding that its induced expression in conventional hESCs is sufficient, alongside naïve-permissive pluripotency conditions, to induce a complete change in phenotype from primed to naïve pluripotency. RNA-seq during the early stages of induction in primed conditions suggests that KLF17 induces this change by regulating genes involved in important signalling pathways. However, we also find that the null mutation of KLF17 in conventional hESCs is not detrimental to naïve resetting or to the growth and survival of the resulting naïve hESCs. Altogether, this suggests that KLF17 functions to regulate genes associated with human naïve pluripotency, but that there is a degree of redundancy *in vitro*, such that KLF17 itself is not strictly necessary for the acquisition and maintenance of naïve hESCs.

## Results

### KLF17 expression in the human embryo is gradually restricted to the epiblast

Detailed single-cell RNA sequencing studies highlight *KLF17* as a molecular marker that is expressed in the human embryonic EPI. First, we reassessed the protein expression dynamics of KLF17 in human embryos, to investigate its distribution across the developing blastocyst. We performed immunofluorescence (IF) analysis of KLF17 alongside the canonical pluripotency factors SOX2 and NANOG in human embryos from the early to late blastocyst stage (FIG.1, S1). NANOG is the earliest known EPI-specific factor in human embryos (Kimber et al., 2008, Niakan and Eggan, 2013), while the *SOX2* expression dynamic closely resembles that of *KLF17* (Blakeley et al., 2015). In keeping with previous data (Kilens et al., 2018), we found that in the earliest stage examined (early day 5 post-fertilisation (dpf)), KLF17 protein was detectable in every cell of the embryo (FIG.1, S1). Though the expression levels across all nuclei were heterogeneous, this widespread staining of KLF17 was largely coincident with SOX2, which is also present in both the inner cell mass (ICM) and trophoectoderm (TE) populations at this stage (FIG.1, S1). As blastocyst development progressed, KLF17 expression was gradually restricted, with ICM enrichment by late day 5 dpf (FIG.1, S1). In early day 7 dpf blastocysts, KLF17 was detectable only in the presumptive EPI cells, delineated by exclusive co-staining with both SOX2 and NANOG (FIG.1, S1). Interestingly, the restriction of KLF17 appeared to progress more slowly than that of SOX2. By late day 5 dpf, SOX2 was only appreciably expressed in the ICM and to a lesser extent in polar TE and it was completely restricted to the EPI in early day 6 dpf embryos (FIG.1, S1). In contrast, there remained appreciable KLF17 protein across cells of the TE in most (3 of 5) of the late day 6 dpf blastocysts analysed (FIG.1, S1). This suggests that the half-life of KLF17 protein may be longer than that of SOX2, given the absence of *KLF17* transcripts in the extraembryonic lineages of human blastocysts by single cell RNA-sequencing (scRNA-seq) analysis (Blakeley et al., 2015). As reported previously, NANOG is detected only in the ICM at all stages of blastocyst development (Niakan and Eggan, 2013). Despite the initial widespread expression pattern of KLF17, its gradual restriction to the NANOG/SOX2 dual-positive EPI suggests that it is specifically retained in the pluripotent EPI, perhaps to perform an unappreciated role in pluripotency regulation or EPI development.

**Figure 1 –.**
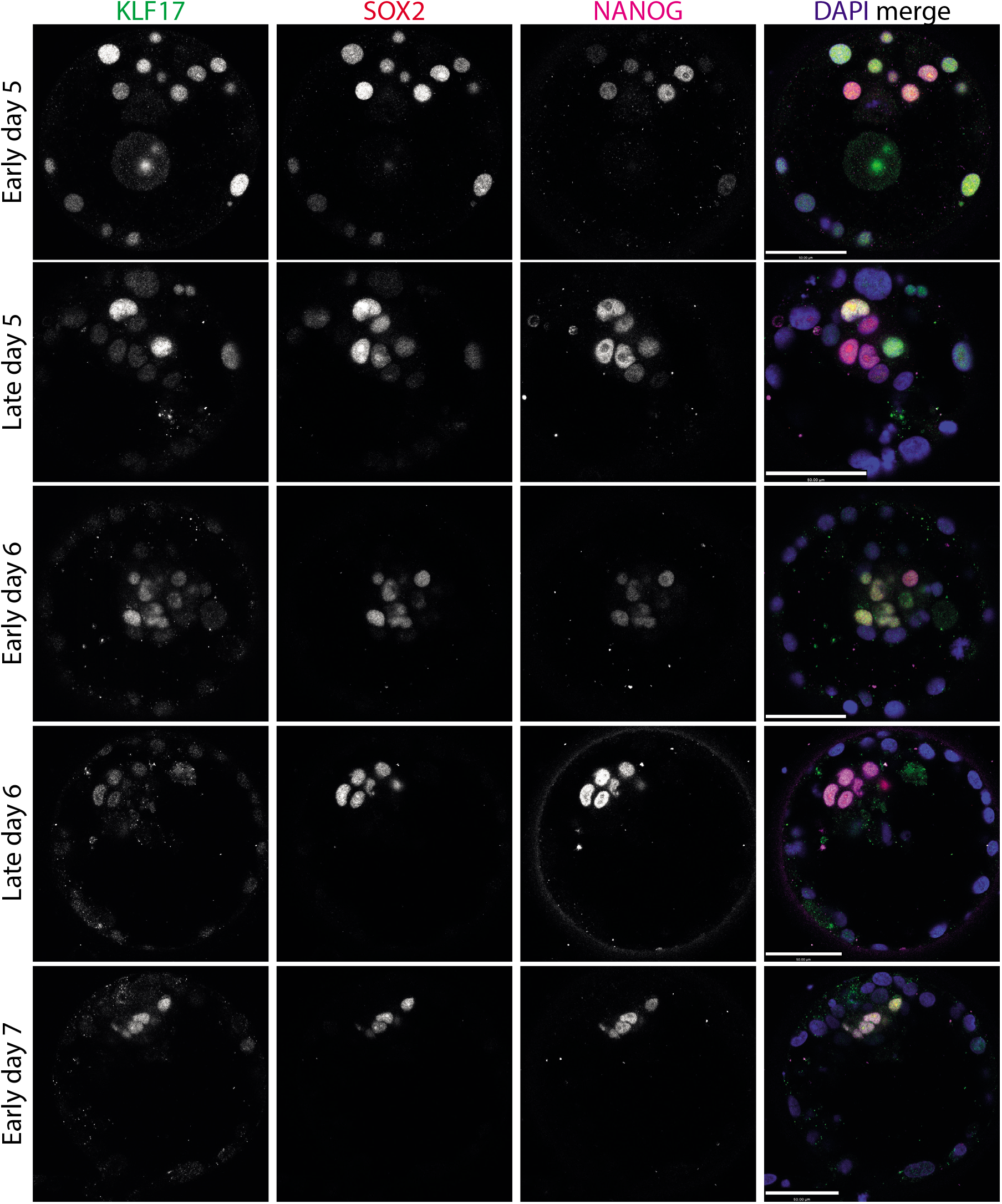
KLF17 expression in the human embryo is coincident with known pluripotency factors. Representative images of immunofluorescence analysis of blastocyst stage human embryos at early day 5 (N = 5), late day 5 (N = 7), early day 6 (N = 9), late day 6 (N = 5) and early day 7 (N = 5) post-fertilisation. Scale bars = 50 μm.

### Induction of KLF17 promotes a naïve pluripotency-like phenotype in conventional hESCs

Given that KLF17 is not expressed when hESCs are cultured in the conventional, primed state, we investigated the effect of ectopic overexpression of *KLF17* in hESCs. We hypothesised that KLF17, as a transcriptional regulator that is enriched in the naïve state (Blakeley et al., 2015, Guo et al., 2016, Messmer et al., 2019), might be sufficient to regulate other naïve hESC-enriched genes when ectopically expressed in primed pluripotent hESCs.

A doxycycline (Dox)-inducible, 3’ HA-tagged *KLF17* transgene was introduced into hESCs by lentiviral transduction (FIG.2A). Initial tests of the transduced line showed that 5 days treatment with 1 μg/ml Dox was sufficient for robust expression of KLF17 protein in almost every cell of the population (FIG.2B). We therefore examined the possibility of gene expression changes in response to ectopic KLF17 in primed culture conditions. Using quantitative reverse transcriptase-polymerase chain reaction (qRT-PCR), we analysed the expression of a number of genes identified as either naïve-or primed-enriched through previous differential gene expression analyses (Stirparo et al., 2018, Messmer et al., 2019) after 5 days Dox induction (FIG.2C). We identified naïve-enriched factors that were significantly upregulated in response to KLF17 induction: *ARGFX* (upregulated ~65-fold average; p~0.03), *REX1/ZFP42* (upregulated ~180-fold average; p~0.02) *DPPA5* (upregulated ~3.9-fold average; p~0.04), *DNMT3L* (upregulated ~300-fold average; p~0.003) and *TFAP2C* (upregulated ~2-fold average; p~0.03). Of these genes, our recent scRNA-seq analysis revealed that under equivalent conditions (H9 cells in mTeSR1 on matrigel) only *REX1/ZFP42* is appreciably expressed in primed hESCs (Wamaitha et al., 2020). Therefore, expression of KLF17 alone is sufficient not only to upregulate a gene already active in conventional hESCs, but also to initiate expression of genes that are otherwise transcriptionally silent. On the other hand, expression of *NANOG* remained transcriptionally unchanged, as did expression of the endogenous *KLF17* gene (FIG.2C).

**Figure 2 –.**
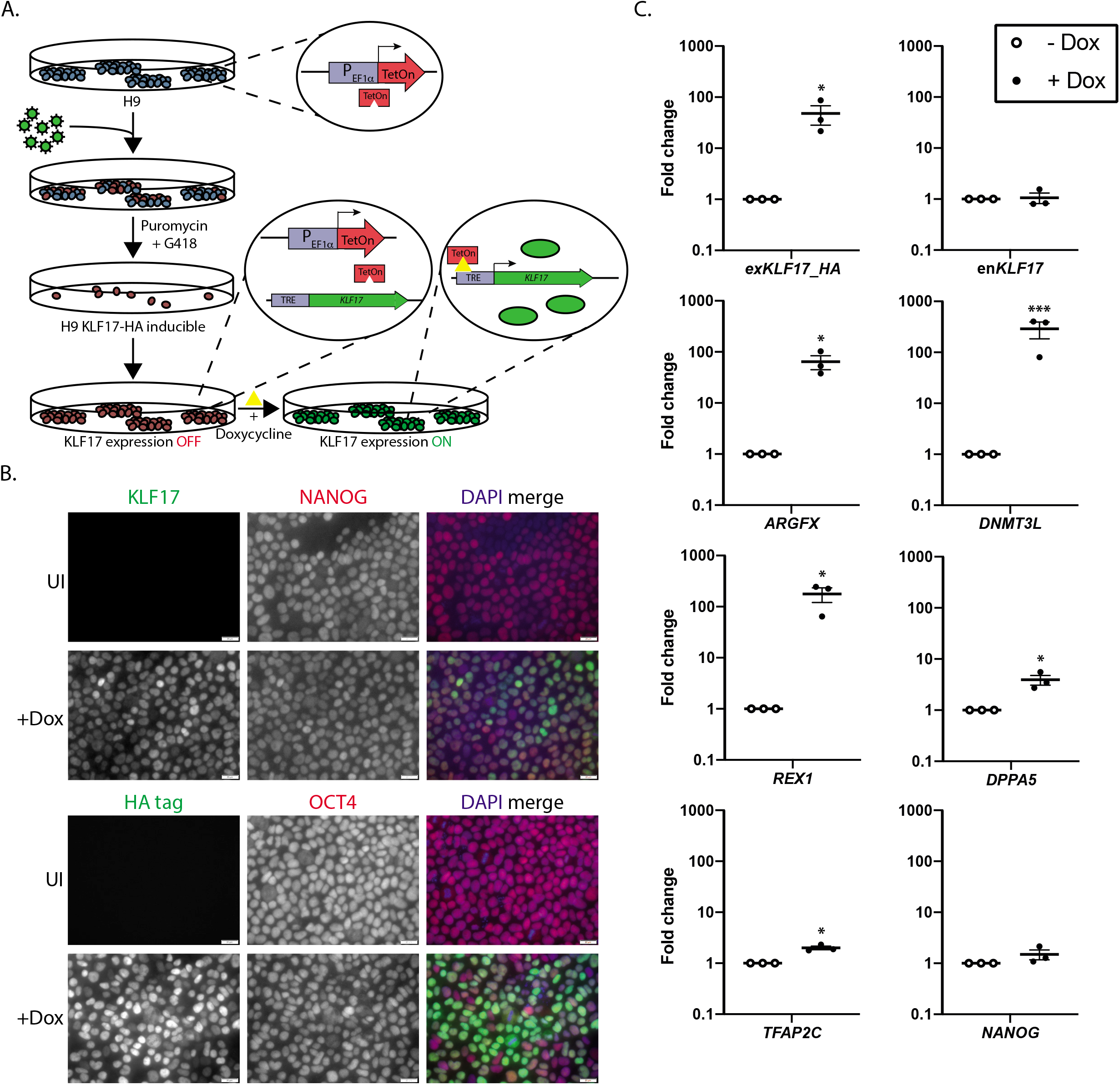
Exogenous KLF17 overexpression induces naïve factor expression in conventional hESCs. **(A)** Schematic diagram of generating H9 KLF17-HA inducible hESCs via lentiviral transduction. **(B)** Immunofluorescence analysis of H9 KLF17-HA inducible hESCs following 5 days uninduced (UI) or 5 days doxycycline (Dox) induction (+Dox). Scale bars = 20 μm. N ≥ 3. **(C)** qRT-PCR analysis of H9 KLF17-HA inducible hESCs following 5 days with (+Dox) or without (-Dox) Dox induction of exogenous KLF17. Relative expression is displayed as fold change versus uninduced cells and normalised to *GAPDH* as a housekeeping gene using the ΔΔCt method. Individual samples are shown as dots, lines represent the mean and whiskers the SEM. Welch’s *t* test, *** *p* < 0.005, * *p* < 0.05.

In order to understand the full extent of the gene expression changes following KLF17 overexpression, we performed mRNA-seq across a 5-day time course of induction. Dimensionality reduction by principal component analysis (PCA) separated the samples by treatment (uninduced (UI) or induced (+Dox)), as well as by timepoint (FIG.3A). Thus, ectopic expression of KLF17 in primed hESCs was sufficient to bring about considerable transcriptome-wide changes.

**Figure 3 –.**
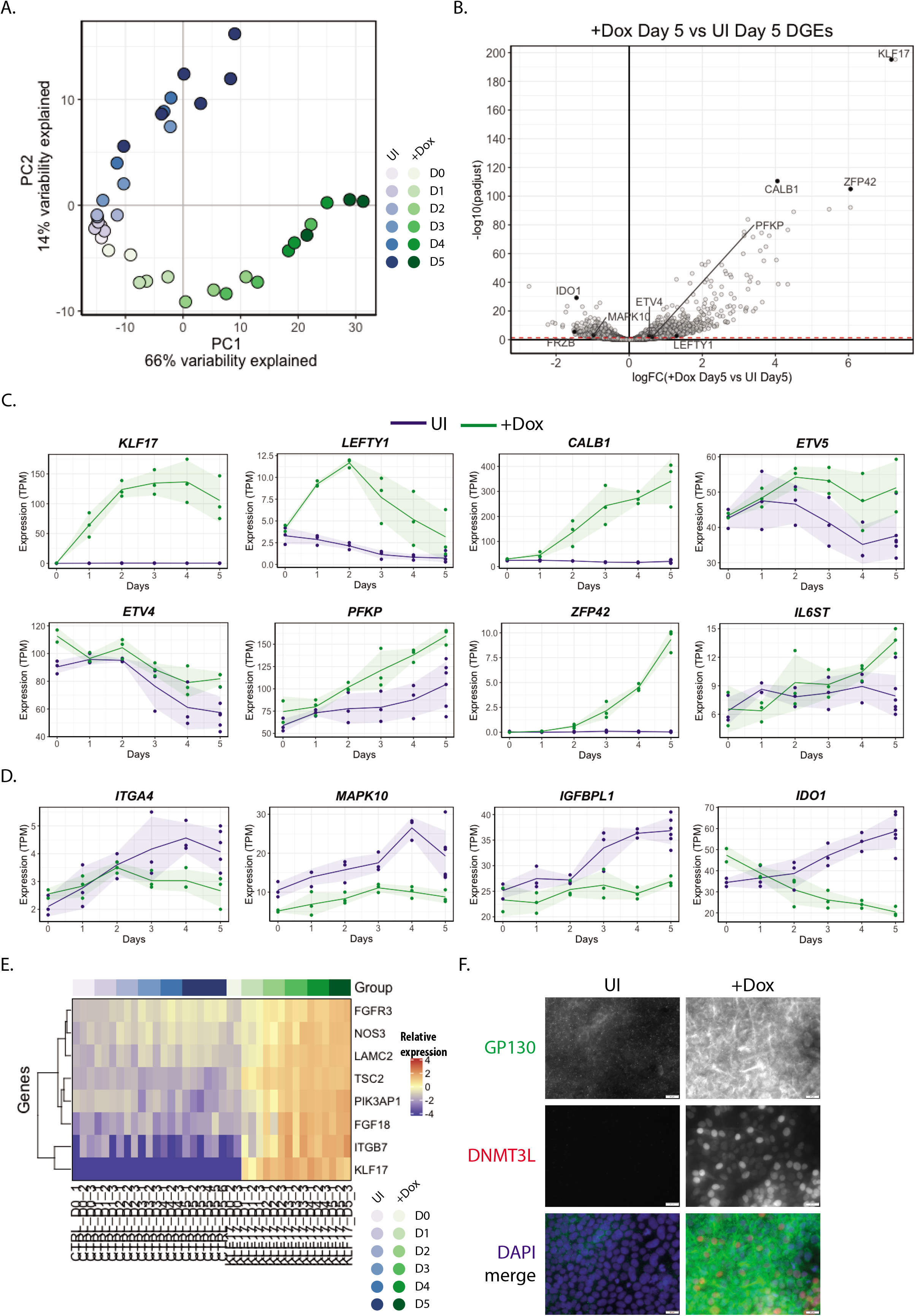
Exogenous KLF17 overexpression induces widespread transcriptional change in conventional hESCs. **(A)** Dimensionality reduction by principal component analysis (PCA) of bulk RNA-seq data collected across a 5-day time course of H9 KLF17-HA inducible hESC growth with (+Dox) or without (UI) Dox induction of exogenous KLF17 expression. **(B)** Volcano plot displaying relative expression of all detected genes in +Dox versus UI H9 KLF17-HA hESCs at day 5 (logFC(+Dox Day5 vs UI Day5)) against the significance of differential expression (-log10(padjust)). The red dotted line notes p_adj_ = 0.05. Individual genes of interest are displayed as filled circles and labelled with the gene name. **(C-D)** Normalised expression (transcripts per million, TPM) of individual genes of interest across the 5-day time course in uninduced control (UI) and KLF17-expressing (+Dox) H9 KLF17-HA hESCs, showing gene that are significantly upregulated (C) or downregulated (D) at day 5. **(E)** Heatmap grouped by sample (UI or +Dox) and time point showing the genes that are highly correlated with *KLF17* across time (Pearson correlation coefficient ≥ 0.85) and fall under the Kyoto Encyclopaedia of Genes and Genomes (KEGG) category “PI3K-Akt signalling pathway”. **(F)** Immunofluorescence analysis of H9 KLF17-HA inducible hESCs following 5 days uninduced (UI) or 5 days doxycycline induction (+Dox). Scale bars = 20 μm. N ≥ 3.

To determine the nature of the genes impacted upon by KLF17, we performed differential gene expression analysis between UI and +Dox samples at each timepoint. At day 5, we uncovered 1760 and 1315 up- and downregulated genes, respectively (p_adj_<0.05) (FIG.3B; Supp Table 1). Intriguingly, of the upregulated genes at day 5, 505 (29%) have been previously identified as enriched in naïve hESCs (Stirparo et al., 2018, Messmer et al., 2019) and/or the human EPI itself (Blakeley et al., 2015), including 46 genes that are EPI-enriched but not differentially expressed between naïve and primed hESCs (e.g. *LEFTY1, CALB1, ETV5, ETV4* and *PFKP*) (FIG.3C). In contrast, of the downregulated genes, 465 (35%) were previously identified as enriched in primed versus naïve hESCs. The negatively regulated, hESC-associated genes included *ITGA2* and *ITGA4, MAPK10, IGFBPL1* and *IDO1* (FIG.3D). This suggests that expression of KLF17 in hESCs cultured under primed culture conditions promotes a shift toward a more naïve pluripotent transcriptome.

It thus appears that KLF17 alone is sufficient to induce significant transcriptional change in primed hESCs over 5 days. To identify those genes most likely to be directly regulated by KLF17, we performed a time course correlation analysis. Using a cut-off for the correlation coefficient of 0.85, we found 70 genes whose expression over time closely mimicked that of *KLF17* itself (FIG.S2A; Supp Table 2). Of these genes, two-thirds (47) were classified as significantly enriched from day 1 (p_adj_<0.05) and almost all (69) were classified as significantly enriched from day 2 onwards (p_adj_<0.05) (FIG.S2B-C), highlighting that these putative KLF17 targets were both rapidly and strongly upregulated following *KLF17* induction. These genes included a number of components of the PI3K-AKT-mTOR signalling pathway (*PIK3AP1, TSC2, NOS3, FGF18, FGFR3, ITGB7* and *LAMC2*; FIG.3E), which we recently showed to be active in both primed and naïve hESCs and a driver of primed hESC and human EPI proliferation (Wamaitha et al., 2020). Following KLF17 overexpression, hESCs significantly upregulated ligands and receptors feeding in to PI3K and the downstream components *NOS3* and *TSC2* (FIG.S3A-G). Of note, all seven such genes are enriched in the human EPI compared to primed hESCs in mTeSR1 on matrigel (Wamaitha et al., 2020), as used in the current experiment. PI3K-AKT-mTOR signalling has also been previously suggested to support an alternative state of naïve pluripotency (Duggal et al., 2015). This suggests that KLF17 induction may modulate signalling through PI3K to a more naïve or EPI-like state.

We investigated this possibility by western blotting, to determine the activation state of the PI3K pathway in UI and +Dox hESCs (FIG.S3Q). We find that phosphorylation of AKT at Serine473 is reduced following 5 days of *KLF17* induction, indicating that the change in expression of PI3K-AKT signalling components in response to KLF17 overexpression is sufficient to adjust the activity of this signalling pathway. In particular, phosphorylation of AKT at Serine473 is usually mediated by the activity of mTOR and stimulates full AKT activity (Alessi et al., 1996, Sarbassov et al., 2005), thereby regulating functions including metabolism, growth and proliferation. A reduction in the levels of Ser473 phosphorylation following upregulation of genes associated with PI3K-AKT-mTOR signalling might indicate negative feedback, acting to keep the *KLF17*-induced hESCs in a signalling steady state.

Other signalling factors were also highly correlated with *KLF17*, including *JAKMIP2, FGFRL1* and *TNFRSF8* and the TGFβ signalling pathway components *LEFTY2* and *TGFB1I1*. A number of cell adhesion-related and cytoskeletal proteins were also included in this list: *LAMC2, MUC4, COL5A1, ITGB7* and *MXRA5* (FIG.S3H-O). It is intriguing that the overexpression of KLF17 alone can influence genes involved in such diverse processes, especially given the importance of changes in morphology and signalling for the conversion of primed to naïve pluripotency. It therefore appears that KLF17 is inducing some of these same resetting-associated changes without any external stimulation.

Of note is the strong expression correlation of *KLF17* and the long non-coding RNA *LINC-ROR* (correlation coefficient 0.921), which is upregulated ~2.4-fold after 24hrs induction (FIG.S3P). *LINC-ROR* has been identified as a regulator of iPSC reprogramming (Loewer et al., 2010) and hESC self-renewal (Wang et al., 2013). *LINC-ROR* expression is regulated by the core pluripotency transcription factors OCT4, SOX2 and NANOG (Loewer et al., 2010) and in turn, acts as a sink for pluripotency destabilising microRNAs (miRNAs) that target the mRNA of these core factors for degradation (Wang et al., 2013). Thus, *LINC-ROR* appears to constitute an important feedback loop for pluripotency maintenance in hESCs. The observed upregulation of *LINC-ROR* expression may therefore protect the KLF17-induced hESCs from differentiation cues, as *LINC-ROR* overexpression has been shown to have a protective effect (Wang et al., 2013).

Finally, we noted that of the 1711 genes downregulated following just 24 hours of Dox induction, there is enrichment for terms related to WNT signalling. Activity of the WNT pathway has been suggested to promote differentiation of hESCs, in both the primed and naïve pluripotent states (Davidson et al., 2012, Singh et al., 2012, Bredenkamp et al., 2019b) and to be suppressed through cross-talk with the PI3K-AKT signalling pathway (Singh et al., 2012). Therefore, downregulation of genes associated with WNT signalling may suggest a further mechanism through which KLF17-overexpressing hESCs would be refractory to differentiation cues. Interestingly, the inability to immediately undergo differentiation in response to established protocols is a feature of naïve hESCs (Rostovskaya et al., 2019).

We confirmed the expression patterns of a number of DEGs by qRT-PCR, as well as validating their upregulation at the protein level, including DNMT3L, VENTX, GP130/IL6ST and TFAP2C, the latter of which is an essential regulator of naïve hESCs (Pastor et al., 2018) (FIG.3F, S4). Altogether this supports our hypothesis that KLF17 acts to transcriptionally regulate genes associated with naïve human pluripotency.

### KLF17 expression drives hESCs to naïve pluripotency alongside signalling modulation

Given that KLF17 is sufficient to upregulate naïve pluripotency-associated factors under conventional primed hESC conditions, we hypothesised that KLF17 induction would be sufficient to reset primed hESCs to a naïve pluripotent state under the appropriate culture regime. The use of ectopic gene expression to drive resetting is common, with deployment of transgenes including *OCT4, KLF4, SOX2, YAP, NANOG* and/or *KLF2* (Hanna et al., 2010, Theunissen et al., 2014, Takashima et al., 2014, Qin et al., 2016), with various media compositions.

Initial testing of KLF17 induction for 5 days under two naïve hESC culture conditions, tt2iL+Gö (Guo et al., 2017) and PXGL (Bredenkamp et al., 2019a, Bredenkamp et al., 2019b), revealed considerably stronger expression of the naïve markers DNMT3L and SUSD2, compared to cells treated equivalently in conventional mTeSR1 medium and untreated controls (FIG.4A, S5A). The upregulation of SUSD2 expression is particularly important to note, as it has been recently identified as a highly specific cell surface marker of naïve hESCs that can be used to enrich for naïve pluripotent cells from a heterogeneous population during resetting (Bredenkamp et al., 2019a). We therefore attempted to propagate the cells, by single cell passaging with MEFs, in both naïve and primed conditions after 5 days of KLF17 induction. Intriguingly, rounded and highly refractile colonies showing typical naïve hESC morphology began to appear only in cells treated with Dox in PXGL medium (FIG.4B, S5B). Conversely, the uninduced cells in PXGL had largely died. In all remaining conditions (mTeSR1 and tt2iL+Gö, +Dox and UI), the *KLF17*-inducible cells survived with either typical primed hESC morphology, that is, flattened, epithelial-like colonies, or evidence of differentiation (data not shown). This suggested that the ectopic expression of KLF17 is sufficient to reset conventional hESCs to a naïve-like pluripotent phenotype when supported by PXGL medium, with modulation of signalling via MEK/ERK (PD0325901), WNT (XAV939, Tankyrase inhibitor), PKC (Gö6983) and LIF/STAT3 (hLIF) (Guo et al., 2017). The unique ability of the PXGL formulation to support this primed-to-naïve transition may reflect its use in the early stages of chemical/epigenetic resetting (Guo et al., 2017), where it appears to perform better than tt2iL+Gö. Additionally, it suggests a particular importance of WNT signalling inhibition (by XAV939), which may relate to the observed downregulation of WNT signalling components in KLF17-induced primed hESCs.

**Figure 4 –.**
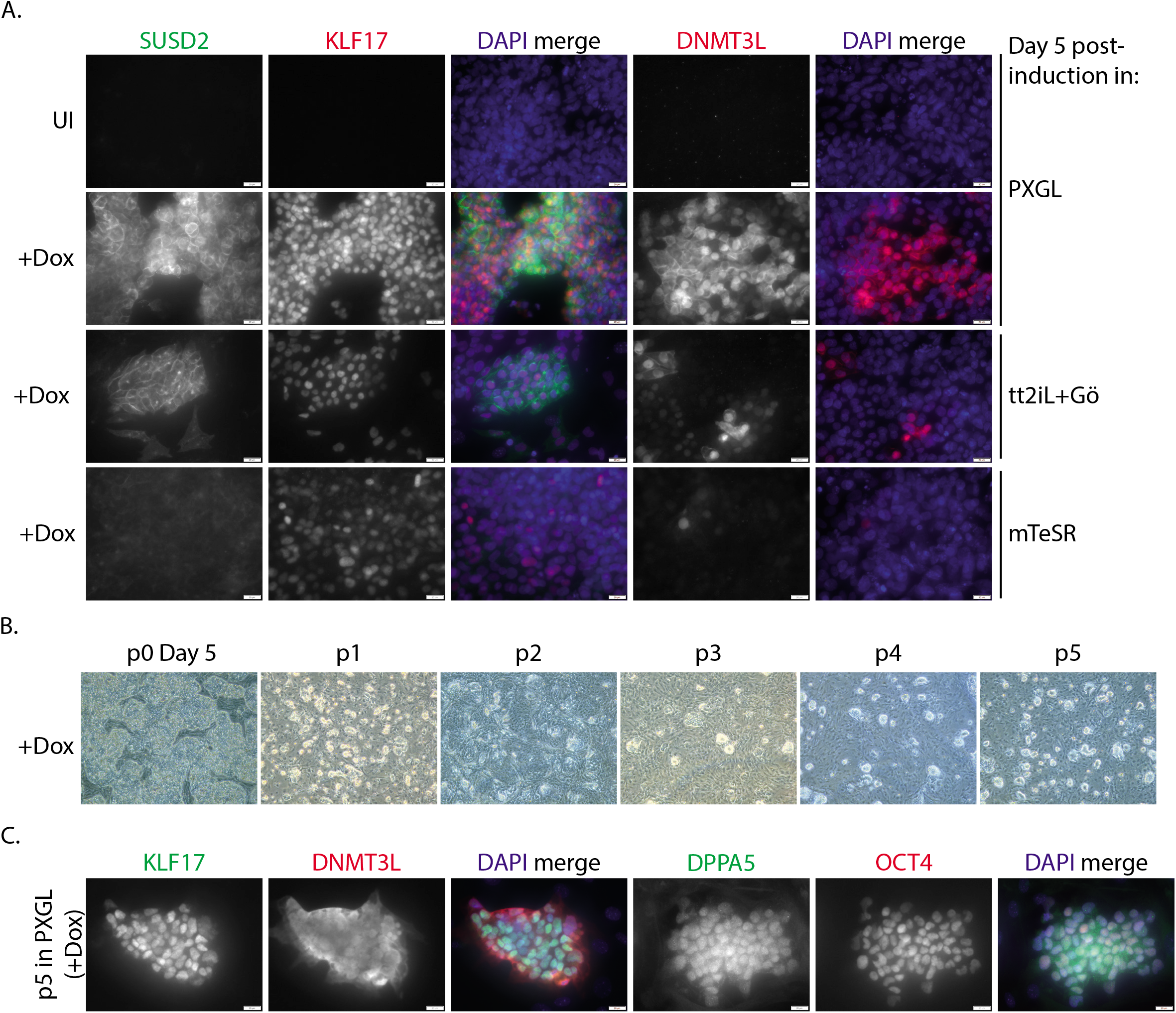
Exogenous KLF17 overexpression is sufficient to drive conventional hESCs to a naïve pluripotent state under PXGL culture. **(A)** Immunofluorescence analysis H9 KLF17-HA inducible hESCs following 5 days uninduced (UI) or 5 days doxycycline induction (+Dox) in the indicated media. Cells were cultured on a mouse embryonic fibroblast (MEF) feeder layer and at 5% O_2_. Scale bars = 20 μm. N ≥ 3. **(B)** Cells induced for 5 days in PXGL medium were uniquely able to give rise to typical naïve hESC-like colonies following serial bulk passaging. **(C)** Representative immunofluorescence analysis of H9 KLF17-HA induced naive hESCs after 4 or 5 passages in PXGL medium. Scale bars = 20 μm. N ≥ 3.

Bulk, single-cell passaging of these naïve-like colonies allowed for stable propagation of *KLF17*-inducible naïve hESCs for a minimum of 5 passages, without requiring additional transgene activation beyond the initial 5-day period of Dox treatment (FIG.4B). We were able to confirm protein expression of a number of naïve hESC markers and factors identified above as upregulated following KLF17 induction in primed culture conditions (FIG.4C). We therefore demonstrated that KLF17 is a potent inducer of the naïve pluripotent state in hESCs.

### Designing a strategy for KLF17 mutation in hESCs

The above investigations revealed that KLF17 is sufficient to induce the naïve pluripotent state in hESCs. Next, we sought to determine whether KLF17 expression is required for resetting of primed hESCs to naïve pluripotency. For this, we designed and optimised a protocol for CRISPR-Cas9-mediated mutation of *KLF17*.

Using *in silico* tools, we designed five guide RNAs (gRNAs) against *KLF17* (FIG.5A, S6A). We considered two strategies to achieve functional knockout of KLF17. First, by targeting the initiating methionine, we aimed to disrupt the entire coding sequence, leading to a complete loss of KLF17 expression. Alternatively, by targeting the functional domain, the DNA-binding ability could be directly disrupted, or a premature termination codon could be introduced, leading to production of a non-functional protein. We introduced Cas9 and each gRNA in turn into primed hESCs and harvested genomic DNA for deep sequencing of the *KLF17* on-target locus by MiSeq analysis. Analysing the proportion of detected insertion and deletion (indel) mutations revealed that the introduction of Cas9 and each of the gRNAs led to indels, with an average mutation efficiency of ~60% (FIG.5B). However, gRNA KLF17(3_3) was clearly inferior and therefore we did not consider it any further.

**Figure 5 –.**
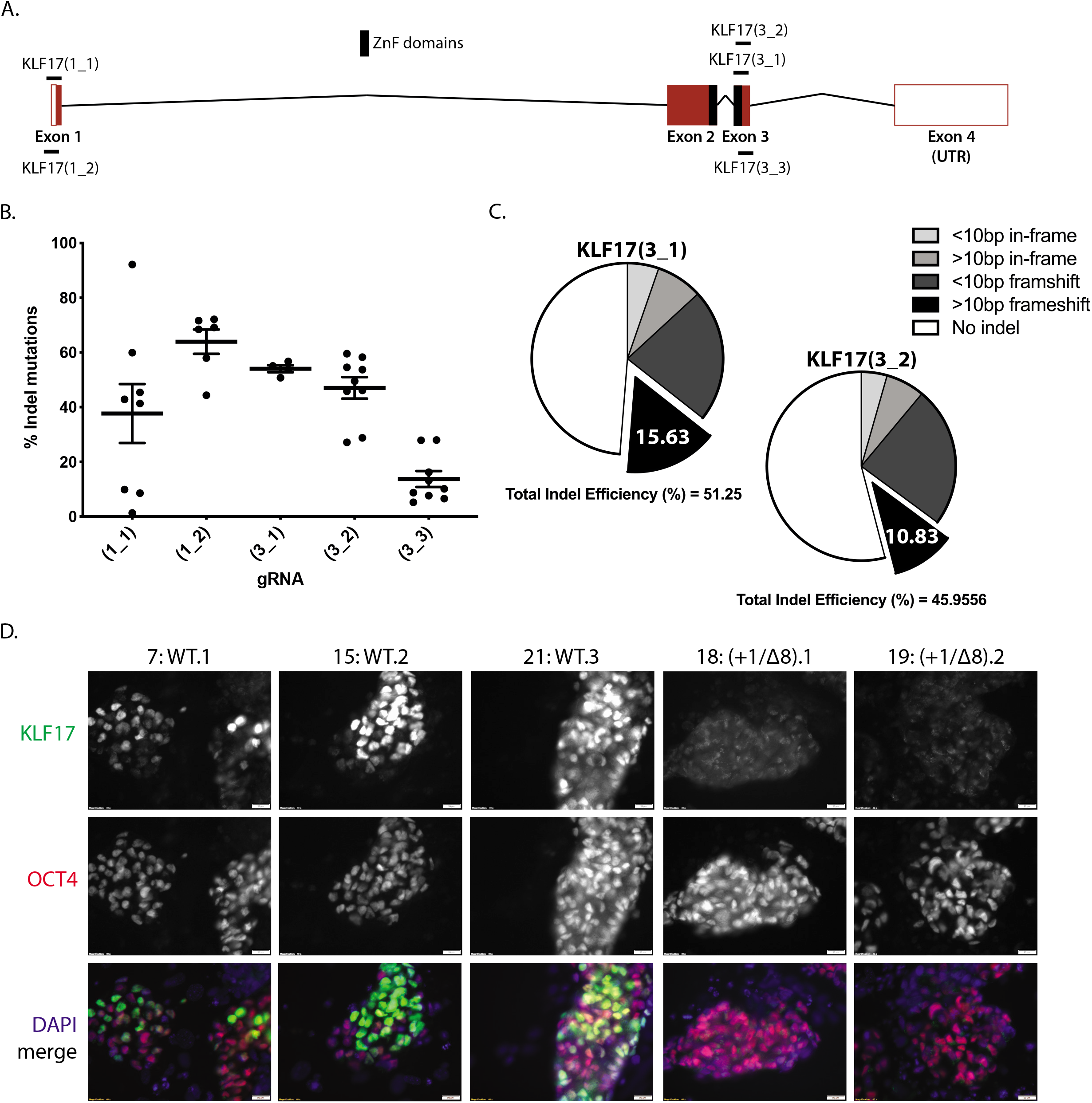
Generating *KLF17-null* mutant hESCs by CRISPR-Cas9. **(A)** Schematic representation of the human *KLF17* locus on Chromosome 1, showing the relative position of the DNA-binding zinc finger domains (filled black rectangles) and the guide RNAs (gRNAs) tested for mutagenic efficiency. Exons are shown as red rectangles, 3’ and 5’ UTR are unfilled rectangles and introns are black chevrons. **(B)** Relative efficiency of each guide shown in (A) measured as a proportion of overall reads containing indel mutations following on-target amplification by MiSeq of the *KLF17* target site. Dots represent individual harvested wells of CRISPR-targeted H9 hESCs, lines represent the mean and whiskers the SEM. **(C)** Pie charts representing the relative proportions of different outcomes of CRISPR-Cas9 editing of H9 hESCs, based on the sequences detected by MiSeq analysis. **(D)** Immunofluorescence analysis of H9 hESCs targeted with Cas9 and gRNA KLF17(3_1) by nucleofection of a plasmid, following subjection to the epigenetic resetting protocol (Guo et al., 2017) for 8 days. Internal wild-type (WT) controls (#7, #15 and #21) are clones that were subjected to nucleofection, puromycin selection and clonal expansion, but were unedited, with a wild-type genotype. Compound null mutant clones (#18 and #19) were verified by MiSeq and IF. N = 3.

In order to decide upon the optimal gRNA for generating KLF17 null mutant (*KLF17^-/-^*) hESCs, we investigated the nature of the indels resulting from CRISPR-Cas9 targeting in each case. Firstly, it was clear that both KLF17(1_1) and KLF17(1_2) were biased towards the introduction of very small indel mutations, with the vast majority of indels less than 10 bp in size (FIG.S6B). Targeting with either of the exon 1-targeted gRNAs would thus leave the possibility of *KLF17* expression from an identified alternative initiating methionine, with the possibility of generating either an essentially wild-type protein or a dominant-negative KLF17 mutant, which could have unexpected consequences. We therefore focussed on the exon 3-targeting gRNAs, KLF17(3_1) and KLF17(3_2). The overall efficiency of these two gRNAs was very similar, but sequence analysis revealed a stronger propensity for the introduction of larger frameshift alleles by KLF17(3_1) (FIG.5C). By disrupting a larger stretch of sequence within the region encoding the KLF17 DNA-binding domain, longer frameshift indels would be expected to be more highly mutagenic. We therefore determined to generate *KLF17^-/-^* hESC lines using gRNA KLF17(3_1).

### KLF17^-/-^ hESCs are not impaired in their ability to adopt naïve pluripotency

Following nucleofection and single-cell amplification of wild-type, primed hESCs, we generated 8 *KLF17*-targeted clones (FIG.S7A). Initial genotyping by short-range PCR and next-generation MiSeq suggested a high proportion of homozygous editing (5 of 8 edited clones; FIG.S7A-B). However, analysis of a ~950 bp region surrounding the on-target site revealed that these apparent homozygous clones had actually undergone an unexpected, long-range editing event on one allele (FIG.S7A-B).

This was apparent from the lack of amplification of both alleles, as determined by the presence of only one variant-type at a highly polymorphic site in the human genome, while the remaining wild-type and heterozygous clones confirmed that this variant is heterozygous in the parental cells (FIG.S7A-C). The extent of the damage was only determined in one clone, #9, where a 163 bp deletion could be detected in the sequence. For the remaining 4 clones, the damage apparently completely prevented amplification of the second allele. This highlights the importance of in-depth genotyping following CRISPR-Cas9-mediated mutagenesis, as previously noted (Kosicki et al., 2018, Cullot et al., 2019, Rayner et al., 2019, Przewrocka et al., 2020, Alanis-Lobato et al., 2020). We therefore sought to test whether clones #18 and #19, compound mutants with two frameshifted alleles predicted to introduce premature stop codons in the sequence encoding the third zinc fingers (FIG.S7D-F), were null for KLF17 expression.

We subjected three wild-type control clones and clones #18 and #19 to chemical resetting (Guo et al., 2017) for eight days and observed robust coexpression of KLF17 and OCT4 in control cells, while the compound mutants lacked detectable KLF17 protein (FIG.5D). To determine whether *KLF17^-/-^* hESCs were able to adopt a naïve pluripotent phenotype, we repeated the chemical resetting and found that both the wild-type controls and *KLF17^-/-^* hESCs could be propagated in tt2iL+Gö conditions for at least 10 passages, maintaining typical naïve morphology (FIG.6A). To identify molecular differences arising in *KLF17^-/-^* hESCs, we performed mRNA-seq at various timepoints throughout the chemical resetting process. This confirmed lack of *KLF17* expression in the compound mutant clones (FIG.6C). The lack of appreciable *KLF17* RNA expression (TPM<5) suggests that the presence of premature termination codons following the CRISPR-Cas9 target site induced nonsense-mediated decay of the mRNA during translation (Nickless et al., 2017). Clones #18 and #19 are therefore *bona fide KLF17-null* mutant hESCs. Despite this, PCA analysis of all samples revealed tight clustering of wild-type and *KLF17^-/-^* hESCs at all timepoints, suggesting that the lack of KLF17 expression did not significantly affect global gene expression (FIG.6B), consistent with the fact that the *KLF17^-/-^* cells were able to reset and survive long-term under naïve culture conditions. This may indicate that KLF17 expression is not required for resetting under the given conditions, or there may be redundancy with other genes that compensate for null mutations in *KLF17*.

**Figure 6 –.**
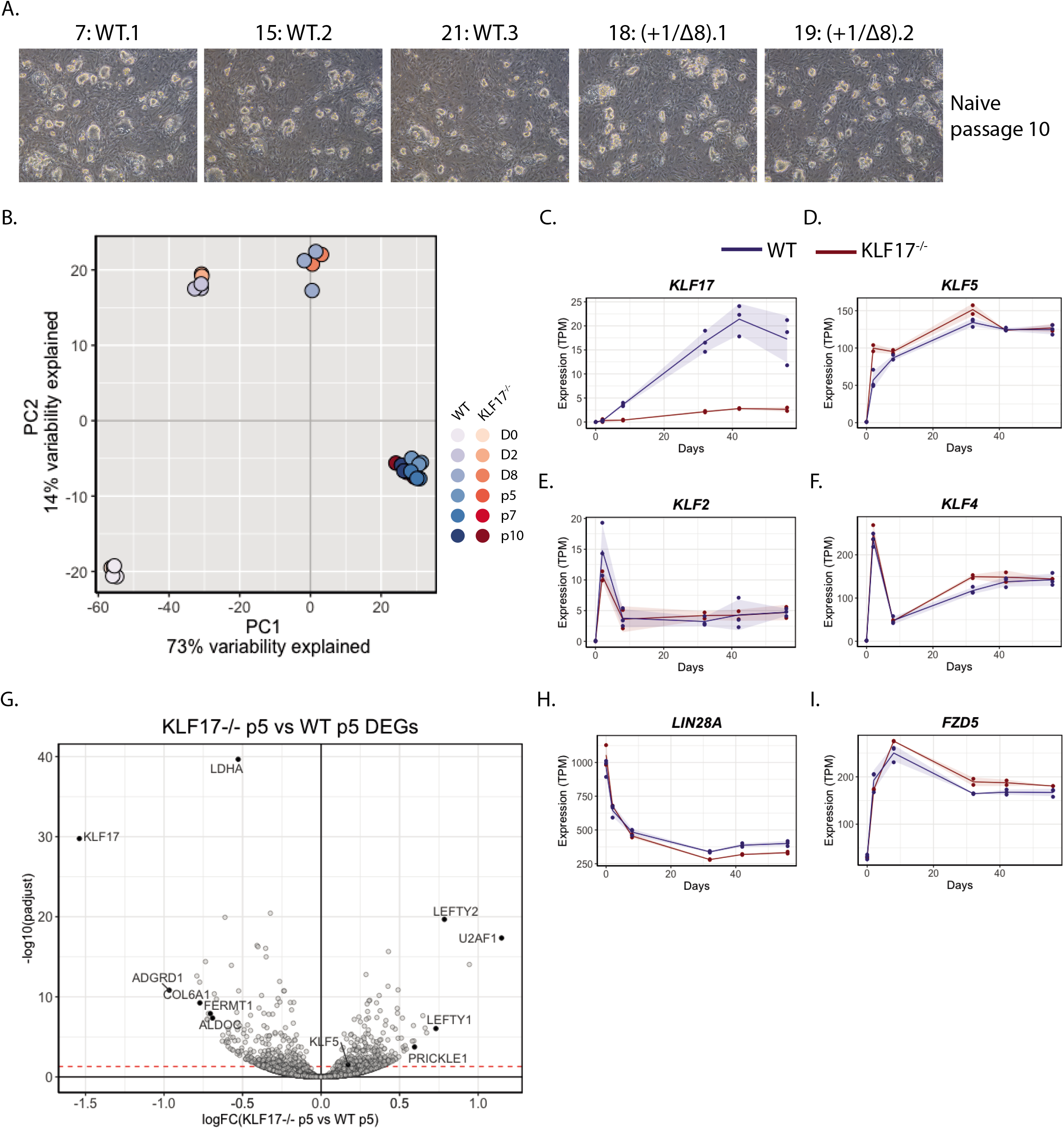
*KLF17*-null hESCs are capable of attaining and maintaining naïve pluripotency. **(A)** Representative brightfield images of WT and *KLF17^-/-^* H9 hESCs following 10 passages under naïve culture conditions. **(B)** Dimensionality reduction by principal component analysis (PCA) of bulk RNA-seq data collected at various times during the epigenetic resetting (Guo et al., 2017) of WT and *KLF17^-/-^* H9 hESCs. **(C-F)** Normalised expression (TPM) of individual genes of interest across the full resetting time course showing (C) lack of appreciable *KLF17* transcripts, (D) temporally-limited upregulation of the paralogue *KLF5* in *KLF17^-/-^* H9 hESCs and (E) equivalent expression of the paralogues *KLF2* and (F) *KLF4*. **(G)** Volcano plot displaying relative expression of all detected genes in *KLF17^-/-^* versus WT naïve H9 hESCs following 5 passages in naïve culture conditions (logFC(*KLF17^-/-^* p5 vs WT p5)) against the significance of differential expression (-log10(padjust)). The red dotted line notes p_adj_ = 0.05. Individual genes of interest are displayed as filled circles and labelled with the gene name. **(H-I)** Normalised expression (TPM) of individual genes of interest across the full resetting time course showing downregulation of (H) the pluripotency-associated factor *LIN28A* and (I) the WNT signalling receptor *FZD5* in *KLF17^-/-^* H9 hESCs.

Interestingly, analysis by DESeq2 identified the *KLF17* paralogue *KLF5* as an early differentially expressed gene, being significantly upregulated in *KLF17^-/-^* versus wild-type naïve hESCs at day 2 of the resetting process (FIG.6D), whereas *KLF2* and *KLF4* are expressed equivalently (FIG.6E-F). It is therefore possible that during the early stage of resetting, where hESCs are undergoing global epigenetic “opening” in response to histone deacetylase inhibition (Guo et al., 2017) and therefore may be in a more phenotypically flexible state than usual, the brief upregulation of *KLF5* is sufficient to compensate for a function carried out by KLF17 in the wild-type state.

Despite this possible redundancy, further analysis revealed a considerable increase in the number of DEGs between wild-type and mutant hESCs at naïve passage 5 (p5), by which point the naïve pluripotent phenotype is suggested to become more stable (Guo et al., 2017). At p5, 316 genes were significantly upregulated and 311 genes were significantly downregulated (p_adj_<0.05) (FIG.6G; Supp Table 3). Enrichment analysis identified significantly downregulated terms related to “Glycolysis/Gluconeogenesis”, “Fructose and mannose metabolism” and “Translation” among others, while upregulated terms included “Signalling by WNT”, “Proteasome” and “Protein processing in endoplasmic reticulum”. Some of the most important, rate limiting enzymes of glycolysis are included in the list of significantly downregulated DEGs, including *HK2, PFKL, ENO1* and *ENO2, PGK1* and *PKM* (FIG.S8A-F). If energy production through glycolytic metabolism is limited, proteasomal degradation might be upregulated as a means to compensate, promoting metabolism of non-essential proteins.

The enrichment terms for downregulated genes at p5 in *KLF17*-null naïve hESCs in fact overlap with those of mRNAs identified as direct targets of the RNA-binding protein LIN28A (Peng et al., 2011), which is itself significantly downregulated from p5 onwards (FIG.6H). LIN28A has been implicated in pluripotency regulation in numerous studies (Heo et al., 2008, Kim et al., 2014, Viswanathan et al., 2008, Yu et al., 2007), though a potential role specifically in naïve hESCs has not been explored. Nevertheless, its significant and maintained downregulation may suggest that its expression depends either directly or indirectly on KLF17 in tt2iL+Gö conditions.

Finally, upregulation of genes associated with WNT signalling in *KLF17^-/-^* naïve hESCs is particularly interesting, given the proposed role of WNT signalling in promoting hESC differentiation (Guo et al., 2017). WNT ligands, receptors and scaffolding proteins are upregulated in the mutant cells (FIG.6I, S8G-H), an effect that might only be observed at such a late timepoint due to the withdrawal of WNT inhibition by XAV939. As stated earlier, KLF17 overexpression in primed hESCs also led to rapid downregulation of a number of WNT components, such that it is an enriched term in the downregulated DEGs at day 1 post-induction (Supp Table 4). This may suggest that WNT signalling is either directly or indirectly regulated by KLF17 transcriptional activity and the continued inhibition of WNT during the early stages of resetting may account for the ability of *KLF17*-null hESCs to adapt to naïve culture.

Altogether this suggests that while KLF5 and KLF17 may display some redundant functions during the establishment of naïve from primed pluripotency in hESCs, *KLF17^-/-^* naïve hESCs are not as phenotypically stable as their wild-type counterparts, since KLF17 function appears to regulate important metabolic and signalling pathways.

## Discussion

In this study, we investigate the human EPI-enriched transcription factor KLF17. By IF analysis of developing human blastocysts, we show that the protein dynamics of KLF17 are remarkably similar to those of the known pluripotency factor SOX2. Both transcription factors display widespread expression across all cells in the early blastocyst, with gradual restriction to the pluripotent EPI, marked by NANOG expression. Therefore, the expression pattern of KLF17 during pre-implantation human development is suggestive of a role in pluripotency regulation.

Indeed, we show that KLF17 is able to induce the expression of a naïve hESC-like transcriptome in primed hESCs and is furthermore sufficient to drive conventional primed to naïve hESC pluripotency. This implies that KLF17 is a powerful inducer of the human naïve pluripotent state *in vitro* and, given its *in vivo* expression, it is interesting to speculate that it may have a role in pluripotency establishment in the human embryo.

From transcriptome analysis of KLF17-overexpressing cells, we suggest that KLF17 may be involved in modulating the expression of components involved in various signalling pathways, primarily the PI3K-AKT and WNT pathways. Of note, the various signalling effectors that are differentially expressed following KLF17 induction are often the most highly correlated with KLF17 expression, as well as being some of the earliest DEGs. In contrast to this, known markers of naïve pluripotency like DNMT3L and SUSD2 are induced much later. This is suggestive of an indirect relationship between KLF17 induction and the expression of naïve pluripotency marker genes, which may well involve the direct action of KLF17 upon signalling effectors to promote naïve pluripotency and inhibit pro-differentiation cues. This is particularly interesting, given the roles of PI3K-AKT (Wamaitha et al., 2020) and WNT signalling in human pluripotency and differentiation (Singh et al., 2012, Bredenkamp et al., 2019b, Mathieu et al., 2019), and the apparent importance of WNT inhibition for recent methods of naïve pluripotency establishment (Zimmerlin et al., 2016, Guo et al., 2017, Bredenkamp et al., 2019b). In naïve hESCs, KLF17 might act to endogenously dampen WNT signalling by impinging upon the expression of components of the WNT pathway, thereby reinforcing the naïve state.

Nevertheless, we also find that loss of KLF17 function is not detrimental to hESC resetting, such that primed *KLF17^-/-^* hESCs are still able to adopt and maintain naïve pluripotency under the conditions investigated herein. This is surprising, given the rapid upregulation of KLF17 expression that has been reported during chemical resetting (Guo et al., 2017) and raises the possibility of genetic compensation. Indeed, *KLF2, KLF4* and *KLF5*, which have all been implicated in pluripotency regulation and to have redundant functions with human *KLF17* in mESCs (Yamane et al., 2018), are also rapidly upregulated in the early stages of resetting in both wild-type and *KLF17^-/-^* hESCs. Furthermore, we observe that *KLF5* is transiently upregulated in *KLF17*-null cells versus wild-type controls at day 2, and again at p5. While *KLF5* transcripts are more abundant in the TE of the human blastocyst (Blakeley et al., 2015), they are still appreciably expressed in both the EPI and PrE. This suggests that *in vitro*, and perhaps *in vivo*, KLF5 may fulfil overlapping functions with KLF17 in the establishment and/or maintenance of naïve pluripotency. Further work could address this question of compensation by performing dual knockout of both *KLF17* and *KLF5* in hESCs and investigating the cells’ competency to undergo chemical resetting. However, the technology for genetic manipulation of human embryos is still in its infancy and so it may unfortunately not be possible to test this hypothesis *in vivo*.

Alternatively, it is feasible that no single factor acts to compensate for KLF17 knockout and instead, a combinatorial action of transcription factors with overlapping targets is able to maintain *KLF17*-null naïve hESCs. This situation would be reminiscent of that observed by manipulating the expression of “peripheral” pluripotency regulators in mouse embryos and mESCs, including KLF2 and KLF4, where individual peripheral factors are mostly dispensable, but together act to reinforce the stability of the pluripotent state mediated by the core factors, OCT4 and SOX2 (Nichols and Smith, 2012). For instance, knockdown of either *Klf2, Klf4* or *Klf5* in naïve mESCs does not appear detrimental (Jiang et al., 2008, Yamane et al., 2018) and *Klf2*-or *Klf4*-null mutant embryos are viable through preimplantation development (Wani et al., 1998, Ehlermann et al., 2003). Despite this, all three factors have validated roles in pluripotency (Parisi et al., 2008, Hall et al., 2009, Jiang et al., 2008).

While *KLF17^-/-^* naïve hESCs did not overtly differ from wild-type counterparts, we did find interesting trends by differential gene expression at p5 of naïve culture, when WNT inhibition by XAV939 is withdrawn. We observe significant downregulation of metabolism and translation, concomitant with upregulation of protein degradation, in *KLF17^-/-^* naïve hESCs. This is reminiscent of the transcriptome changes observed in primed hESCs treated with proteasome inhibition (Saez et al., 2018), which induces cells to enter a state of stress, reflected in their transcriptome. Thus, this overlap of the response of *KLF17^-/-^* hESCs during naïve resetting and proteasome inhibition in primed cells may indicate an induction of the cellular stress response. In particular, the transcriptome changes are suggestive of metabolic stress at a key point during the stabilisation of the naïve pluripotent state, as we observe downregulation of key glycolytic enzymes. This may imply that without exogenous WNT inhibition to stabilise the naïve pluripotent state, and indeed with the observed upregulation of WNT signalling components, KLF17-null naïve hESCs become temporarily unstable.

This could also be reflected in the observed and consistent downregulation of LIN28A, as LIN28A has been directly implicated in the growth and survival of hESCs (Peng et al., 2011). By binding RNA, LIN28A regulates expression primarily at the post-transcriptional level. Thus, there may be further effects in *KLF17^-/-^* naïve hESCs at the protein level, not reflected in the current transcriptome analysis. At p5, *LIN28B* is also significantly downregulated. Interestingly, LIN28 has been recently identified as a naïve-specific marker in porcine ESCs (Chen et al., 2020), at both the RNA and protein level, suggesting that its downregulation may be somewhat detrimental also in naïve hESCs.

Overall, our overexpression studies show that KLF17 may typically have a role in regulating naïve pluripotency *in vitro*. However, it is clear from our data that KLF17 expression is not necessary for establishing naïve hESCs, although our current work does not exclude the possibility that established naïve hESCs are sensitive to loss of KLF17, as recently suggested (Bayerl et al., 2020). We theorise that in a wild-type situation KLF17 may act as a peripheral pluripotency factor in human naïve pluripotency, acting alongside a core pluripotency network of OCT4 and SOX2 to maintain robustness of the pluripotent state and help to limit premature differentiation. This is borne out by its downregulation of the differentiation-promoting WNT signalling pathway and its upregulation of factors like LINC-ROR, which are known to limit differentiation.

However, we also note that the lack of KLF17 necessity in naïve hESC establishment does not rule out a more central role in pluripotency in the human embryo. To date, there have been no systematic comparisons of the outcomes of specific gene modulation in naïve hESCs versus the human pluripotent epiblast, but evidence suggests that they would not necessarily be conserved. For instance, *Nanog-null* naïve mESCs, while prone to differentiation, are still functionally pluripotent, with the capability for chimaera formation (Chambers et al., 2007). In contrast, a *Nanog-null* mouse embryo is unable to form a functional blastocyst or continue development from the peri-implantation stage onward (Mitsui et al., 2003, Chambers et al., 2003, Messerschmidt and Kemler, 2010, Frankenberg et al., 2011). This shows that the *in vivo* phenotype of genetic knockouts can be more severe than that observed *in vitro*. Furthermore, while knockdown of *POU5F1* in hESCs causes the expected differentiation phenotype (Wang et al., 2012), even partial loss of OCT4 function in the human embryo had a much more drastic phenotype, with non-cell-autonomous effects across all three lineages at the blastocyst stage (Fogarty et al., 2017). For this reason, future investigation of the function of KLF17 in human *in vivo* pluripotency is an important next step.

## Materials and methods

### Human embryo thaw and culture conditions

Human embryos at various developmental stages that were surplus to family building desires were donated to the Francis Crick Institute for use in research projects under the UK Human Fertilisation and Embryology Authority License number R0162. Slow-frozen blastocysts (day 5 and day 6) were thawed using the BlastThaw (Origio; 10542010A) kit using the manufacturer’s instructions. Vitrified blastocysts (day 5 and day 6) were thawed using the vitrification thaw kit (Irvine Scientific; 90137-SO) following the manufacturer’s instructions. Human embryos were cultured in pre-equilibriated Global Media (Life Global) supplemented with 5 mg/ml Life Global HSA (LifeGlobal; LGPS-605) and overlaid with mineral oil (Origio; ART-4008-5P and incubated in Embryoscope+ time lapse incubator (Vitrolife).

### Maintenance of standard hESC cultures

Human embryonic stem cells (hESCs) were routinely cultured in mTeSR1 medium (Stem Cell Technologies) on growth factor-reduced Matrigel-coated dishes (BD Biosciences) and passaged as clumps at ~1:20 ratio using ReLeSR (Stem Cell Technologies). Cells were maintained in humidified incubators at 37°C, 5% CO_2_.

### Naïve hESC culture

All naïve hESCs were cultured under hypoxia (5% O_2_, 5% CO_2_), according to recently published protocols (Guo et al., 2017, Bredenkamp et al., 2019b) on mitotically-inactivated DR4 MEFs (prepared in-house) plated at a density of 1×10^6^ per well of a 6-well plate 12-16hrs prior to hESC seeding. Naïve hESCs were passaged as single cells by 4 minutes treatment with Accutase (Thermo Fisher) at 37°C, at split ratios between 1:3 and 1:6, every 3 to 6 days. For culture in (t)t2iL+Gö, 10 μM ROCK inhibitor (Y-27632, Tocris Bioscience) was added overnight before and after passaging, to aid survival. In-house generated, chemically reset, naïve H9 cells were maintained in tt2iL+Gö, with 0.3 μM CHIR99021 (Guo et al., 2017), and XAV supplementation until naïve passage 5.

### Generation and culture of overexpression hESC lines

Doxycycline-inducible overexpression of HA-tagged proteins was achieved using the Lenti-X Tet-On 3G Inducible Expression System (Clontech) following the manufacturer’s protocol, and as outlined previously (Wamaitha et al., 2015). Lentiviral packaging was achieved using 7 μg of transgene-containing plasmid and the Lenti-X Packaging Single Shot reagents. Lentiviral supernatant was harvested after 48hrs and concentrated by ultracentrifugation. To produce stably-transduced cells, hESCs were plated under standard conditions and changed into fresh medium the following morning. 24hrs post-plating, 10 μl concentrated virus was added to hESCs for transduction overnight (~16hrs). hESCs were dual selected with 150 μg/ml G418 and 0.5 μg/ml puromycin 48hrs post-transduction. For induction of transgene expression, doxycycline was added to mTeSR1 medium at 1 μg/ml. For the RNA-seq experiments, *KLF17*-inducible hESCs were plated as normal and induction initiated after 24 hours by addition of 1 μg/ml Dox to the culture medium (mTeSR1). At ~30 hours, a day 0 (pre-induction) control sample was collected, then both induced (+Dox) and uninduced (UI) samples were collected at 24hr intervals from 48hrs (day 1 post-induction) until 144hrs (day 5 post-induction). RNA was extracted from the samples and subjected to bulk RNA sequencing.

### KLF17-driven resetting of primed to naïve hESCs

H9 KLF17-HA inducible hESCs were pre-treated overnight with 10 μM ROCKi, then harvested from standard culture (mTeSR1 on matrigel) by 5 minutes incubation at 37°C with accutase, resuspended in culture medium supplemented with 10 μM ROCKi and counted. 2×10^5^ hESCs were plated per 6-well pre-coated in DR4 MEFs and the cells placed at hypoxia (5% O_2_, 5% CO_2_) for ~24hrs. The following day (day 0), medium was changed to PXGL supplemented with 1 μg/ml Dox. From day 2, medium was replenished each day with PXGL freshly supplemented with 1 μg/ml Dox. On day 5, cells were passaged by 4min incubation in accutase and plated in PXGL with 10 μM ROCKi at a split ratio between 1:5 and 1:20, dependent upon density. Within 24hrs, the cells adopted a domed morphology with highly-refractile colony edges. Cells were passaged again on day 7 or 8 and could subsequently be maintained similarly to chemically-reset cells (Guo et al., 2017), with passaging every 3-4 days at split ratios of between 1:3 and 1:6.

### Design of gRNAs

Guide RNAs (gRNAs) were designed in a non-biased manner against the whole cDNA sequence using a standard design tool (Hsu et al., 2013). For initial screening, gRNAs were selected on the following criteria: (i) *in silico* score is ≥60; (ii) identified off-target sites have ≥3 mismatches; (iii) there are no (or very low frequency, ≤0.1%) single nucleotide polymorphisms (SNPs) occurring in the target sequence; (iv) the gRNA target site falls across an annotated DNA-binding domain.

### Transient nucleofection of hESCs

For cell line testing of CRISPR-Cas9 efficiency, gRNAs were individually cloned into pSpCas9(BB)-2A-Puro (PX459) V2.0 (Addgene plasmid #62988) (Ran et al., 2013), using the BbsI restriction sites. Nucleofection was carried out on an Amaxa 4D-Nucleofector (Lonza) with 4 μg plasmid. 24 hours prior to nucleofection, H9 hESCs were treated with 10 μM Y-27632 (Tocris Bioscience). hESCs were harvested as single cells by Accutase treatment (5min, 37°C) and counted with an automatic cell counter (Nucleocounter NC-200, ChemoMetec). For each gRNA, 2×10^6^ cells were resuspended in 100 μl P3 Primary Cell 4D-Nucleofector X Solution and transferred to nucleocuvettes with 4 μg plasmid. Nucleofection was performed with the pre-set H9 hESC programme (CB-150), then cells resuspended in antibiotic-free mTeSR1 medium supplemented with 10 μM Y-27632 and plated across half of a 6-well plate coated with DR4 MEFs to aid attachment and survival. After 24hrs, medium was changed to mTeSR1 supplemented with 0.5 μg/ml puromycin for 48hrs. Cells were allowed to recover for 8 days prior to harvesting for DNA extraction and assessment of CRISPR-Cas9 editing efficiency by MiSeq analysis. On-target editing was assessed by next-generation sequencing on the MiSeq platform (Illumina), and editing efficiency determined by analysing the FastQ files using both the Cas-Analyzer tool from CRISPR RGEN Tools (Park et al., 2017) and the CrispRVariants package in R (Lindsay et al., 2016).

### Generation of clonal knockout hESCs

H9 hESCs were first nucleofected with 4 μg pSpCas9(BB)-2A-Puro (PX459) V2.0 containing the gRNA KLF17(3_1) as described above. Following 48hrs treatment with 0.5 μg/ml puromycin, cells were allowed to recover on DR4 MEFs for ~10 days, then manually passaged as single cells following treatment with Accutase (5 minutes, 37°C) or Accumax (10 minutes, 37°C; Sigma Aldrich) at clonal density into Matrigel-coated 24-well tissue culture plates (Corning). Cells were sub-cloned once more by manual picking and single-cell dissociation into 12-well plates, then 24 clones passaged in duplicate and assessed for KLF17 mutation by on-target Sanger sequencing and MiSeq analysis.

### Immunofluorescence analysis

Cultured cells were fixed with 4% paraformaldehyde (PFA) in PBS for 1hr at 4°C, then permeabilised in PBS containing 0.5% Tween-20 (PBS-T(0.5%)) for 20 minutes at room temperature. Blocking was carried out for 1hr at RT in PBS-T(0.1%) with 10% donkey serum. Primary antibodies were diluted as listed in TABLE 1 in blocking solution, and incubated overnight at 4°C. Cells were washed several times in PBS-T(0.1%), then incubated in secondary antibodies in blocking solution for 1hr at RT. Following repeated washing, cells were treated with DAPI-Vectashield mounting medium (Vector Labs) at 1 in 30 in PBS-T(0.1%), prior to imaging on an Olympus IX73 (Olympus Corporation).

**Table 1 -.** Primary and secondary antibodies used in immunofluorescence

| Target | Species | Dilution | Supplier | Catalogue Number |
| --- | --- | --- | --- | --- |
| Anti-DNMT3L | Mouse | 1 in 500 | Abcam | ab93613 |
| Anti-GP130 | Rabbit | 1 in 250 | Thermo Fisher | PA5-80735 |
| Anti-HA (3F10) | Rat | 1 in 500 | Sigma Aldrich (Roche) | 11867423001 |
| Anti-KLF17 | Rabbit | 1 in 500 (hESCs)<br>1 in 200 (embryo) | Atlas Antibodies | HPA024629 |
| Anti-NANOG | Goat | 1 in 200 | R&D Systems | AF1997 |
| Anti-OCT4 | Mouse | 1 in 100 | Santa Cruz Biotechnology | SC-5279 |
| Anti-SOX2 | Rat | 1 in 100 | Invitrogen | 14-9811-82 |
| Anti-SUSD2 | Mouse | 1 in 250 | Biolegend | 327401 |
| Anti-TFAP2C | Goat | 1 in 200 | R&D Systems | AF5059 |
| Anti-VENTX | Rabbit | 1 in 500 | Cambridge Bioscience | HPA050955 |
| Alexa Fluor anti-mouse IgG | Donkey | 1 in 300 | Invitrogen | A21202 (488 nm)<br>A21203 (594 nm)<br>A31571 (647 nm) |
| Alexa Fluor anti-rabbit IgG | Donkey | 1 in 300 | Invitrogen | A21206 (488 nm)<br>A21207 (594 nm)<br>A31573 (647 nm) |
| Alexa Fluor anti-goat IgG | Donkey | 1 in 300 | Invitrogen | A11055 (488 nm)<br>A11058 (594 nm)<br>A21447 (647 nm) |
| Alexa Fluor anti-rat IgG | Donkey | 1 in 300 | Invitrogen | A21208 (488 nm)<br>A21209 (594 nm) |

For human embryos, fixation was performed in 4% PFA in PBS for 1hr at 4°C, then the embryos permeabilised in PBS containing 0.5% Triton-X100 (PBS-Tx(0.5%)) for 20 minutes at RT. Blocking was performed for 1hr at RT in PBS-Tx(0.2%) containing 10% donkey serum and 3% bovine serum albumin (BSA). Primary antibodies were diluted as listed in TABLE 1 in blocking solution, and incubated overnight at 4°C. Embryos were washed several times, then incubated in secondary antibodies in blocking solution for 1hr at RT. Following repeated washing, embryos were transferred into DAPI-Vectashield mounting medium (Vector Labs) at 1 in 30 in PBS-T(0.1%) on coverslip dishes (MatTek), and imaged on a Leica SP8 inverted confocal microscope (Leica Microsystems).

### RNA isolation from hESCs and qRT-PCR

RNA was isolated using TRI reagent (Sigma) and DNase I-treated (Ambion). cDNA was synthesized using a Maxima first strand cDNA synthesis kit (Fermentas). qRT-PCR was performed using SensiMix SYBR low-ROX kit (Bioline) on a QuantStudio5 machine (Thermo Fisher). Primers pairs used are listed in TABLE 2. Each sample was run in triplicate. Gene expression was normalized using *GAPDH* as the housekeeping gene, and the results analysed using the ΔΔCt method.

**Table 2 –.**
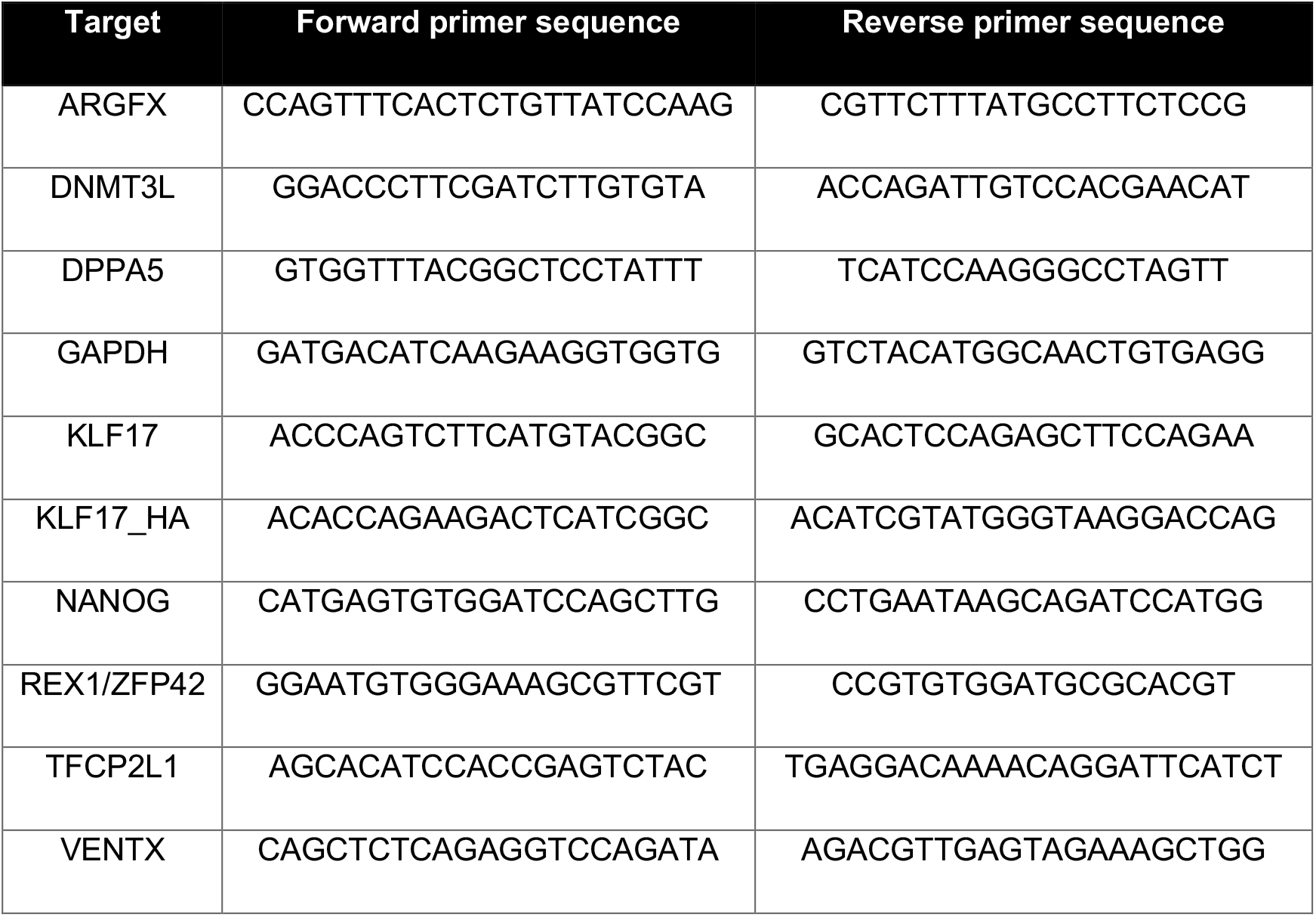
Primers used for qRT-PCR

### RNA sequencing

For RNA-seq, RNA was isolated and DNase-treated as above, and libraries were prepared using the polyA KAPA mRNA HyperPrep Kit (Roche). Quality of submitted RNA samples and the resulting cDNA libraries was determined by ScreenTape Assay on a 4200 TapeStation (Agilent). Prepared libraries were submitted for single-ended 75 bp sequencing on an Illumina HiSeq 4000 (Illumina).

### Genomic DNA extraction

Total genomic DNA was extracted from hESCs using the DNeasy Blood and Tissue Kit (Qiagen) following manufacturer’s instructions. The concentration and purity of extracted DNA was measured using a nanodrop (DeNovix).

### Protein extraction and quantification

hESCs were harvested for protein extraction by addition of CelLytic M lysis buffer (Merck), freshly supplemented with protease inhibitors (PIC, cOmplete, EDTA-free protease inhibitor cocktail, Roche) and phosphatase inhibitors (PhIC, phosSTOP phosphatase inhibitor, Roche), directly onto plated cells. Cells were scraped, then incubated in lysis buffer for 15min at 4°C. The lysate was collected and clarified by centrifugation at 20,000xg for 15min at 4°C. Protein concentration in the lysates was determined using the BCA assay, then proteins denatured by addition of 4x Laemmli sample buffer (Thermo Fisher) and heating at 90°C for 5 minutes.

### Protein detection by western blotting

Denatured proteins were thawed at 65°C for 5 minutes and vortexed to ensure homogeneity. 20 μg protein per lane was loaded onto 10% Mini-PROTEAN TGX Stain-free protein gels (BIORAD), alongside 5 μl PageRuler Prestained Protein Ladder (Thermo Scientific), and electrophoresed at 100-200 V for one to two hours in a Mini-PROTEAN Tetra Vertical Electrophoresis Cell (BIORAD). Proteins were transferred onto PVDF membranes (TransBlot Turbo Mini PVDF Transfer Packs, BIORAD) using a Trans-Blot Turbo Transfer System (BIORAD). PVDF membranes were blocked for one hour in TBS-T(0.1%) containing 5% non-fat milk and incubated with primary antibodies diluted in either 5% milk or 5% BSA in TBST-T(0.1%) as shown in TABLE 3 overnight at 4°C. Following washes with TBS-T(0.1%), membranes were incubated with secondary antibodies in 5% milk for one hour at room temperature. Proteins of interest were visualised using the SuperSignal West Dura Extended Duration Substrate or SuperSignal West Femto Maximum Sensitivity Substrate (Thermo Scientific) and imaged on an Amersham Imager 600RGB (GE Healthcare).

**Table 3 -.** Primary and secondary antibodies used in western blot

| Target | Species | Dilution | Supplier | Catalogue Number |
| --- | --- | --- | --- | --- |
| Anti-alpha tubulin | Mouse | 1 in 1000 | Sigma Aldrich | T9026 |
| Anti-pan AKT | Rabbit | 1 in 1000 | Cell Signaling Technologies | 9272 |
| Anti-phospho AKT (Ser473) | Rabbit | 1 in 1000 | Cell Signaling Technologies | 9271 |
| Anti-KLF17 | Rabbit | 1 in 500 | Atlas Antibodies | HPA024629 |
| Anti-pan S6 | Mouse | 1 in 1000 | Cell Signaling Technologies | (54D2) #2317 |
| Anti-phospho S6 | Rabbit | 1 in 1000 | Cell Signaling Technologies | #2211 |
| Anti-mouse IgG (H+L), HRP-conjugated | Goat | 1 in 20000 | Cell Signaling Technologies | 7076 |
| Anti-rabbit IgG (H+L), HRP-conjugated | Goat | 1 in 20000 | Cell Signaling Technologies | 7074 |
| Anti-goat IgG (H+L), HRP-conjugated | Donkey | 1 in 20000 | Santa Cruz Biotechnology | SC-2020 |
| Anti-rat IgG (H+L), HRP-conjugated | Goat | 1 in 20000 | Cell Signaling Technologies | 7077 |

### Bulk RNA-sequencing analysis

The ‘Trim Galore!’ utility version 0.4.2 was used to remove sequencing adaptors and to quality trim individual reads with the q-parameter set to 20 (https://www.bioinformatics.babraham.ac.uk/projects/trim_galore/ (retrieved 03-05-2017)). Then sequencing reads were aligned to the human genome and transcriptome (Ensembl GRCh38 release-89) using RSEM version 1.3.0 (Li and Dewey, 2011) in conjunction with the STAR aligner version 2.5.2 (Dobin et al., 2013). Sequencing quality of individual samples was assessed using FASTQC version 0.11.5 (https://www.bioinformatics.babraham.ac.uk/projects/fastqc/ (retrieved 03-05-2017)) and RNA-SeQC version 1.1.8 (DeLuca et al., 2012). Differential gene expression was determined using the R-bioconductor package DESeq2 version 1.24.0 (Love et al., 2014). Within the DESeq2 package, adjusted p values for log-fold changes were calculated using the Benjamini-Hochberg method and the betaPrior parameter was set to “TRUE”. For the *KLF17^-/-^* hESCs in naïve conditions, each timepoint was normalised individually, to account for the significant cell-state changes occurring across the long time course of the experiment (~60 days).

## Acknowledgements

We thank the donors of human embryos whose contributions enable this research. Further, we thank all members of the Niakan lab for their technical assistance and for help and comments on the paper, and James Briscoe, Robin Lovell-Badge and Jonathan Chubb for advice throughout the project. We are grateful to the Francis Crick Institute’s Science Technology Platforms: Robert Goldstone and Deb Jackson from the Advanced Sequencing Facility; Advanced Light Microscopy; and the Genomics Equipment Park.

## Author Contributions

R.A.L. performed the experiments with assistance from A.M. S.B. performed major bioinformatics analysis with assistance from R.A.L. K.K.N. and R.A.L. conceived the study and analysed data. R.A.L. wrote the manuscript with feedback from all authors.

---

**Figure S1 –.**
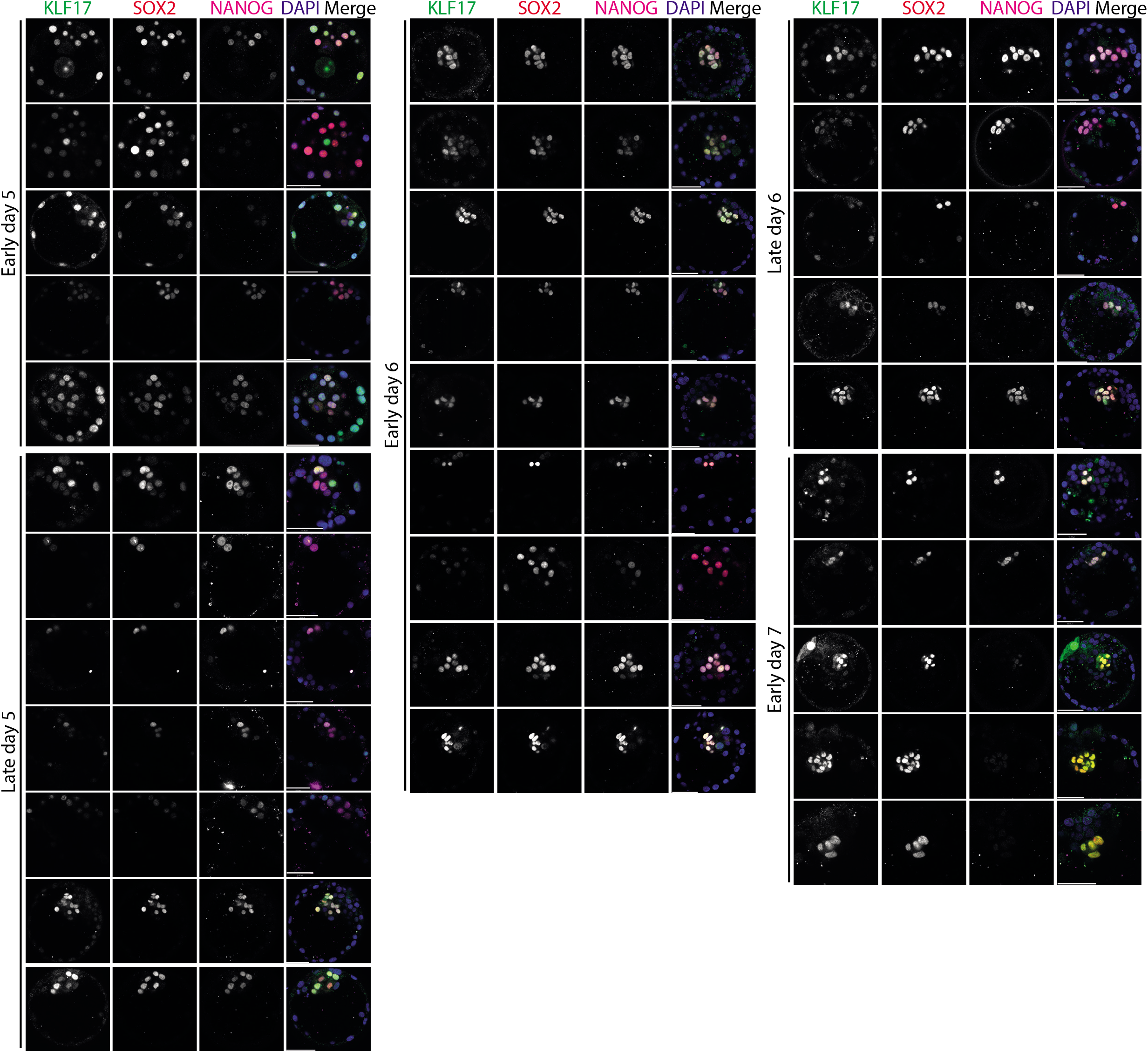
KLF17 expression in the human embryo is coincident with known pluripotency factors. Immunofluorescence analysis of blastocyst stage human embryos at early day 5 (N = 5), late day 5 (N = 7), early day 6 (N = 9), late day 6 (N = 5) and early day 7 (N = 5) post-fertilisation. Scale bars = 50 μm.

**Figure S2 –.**
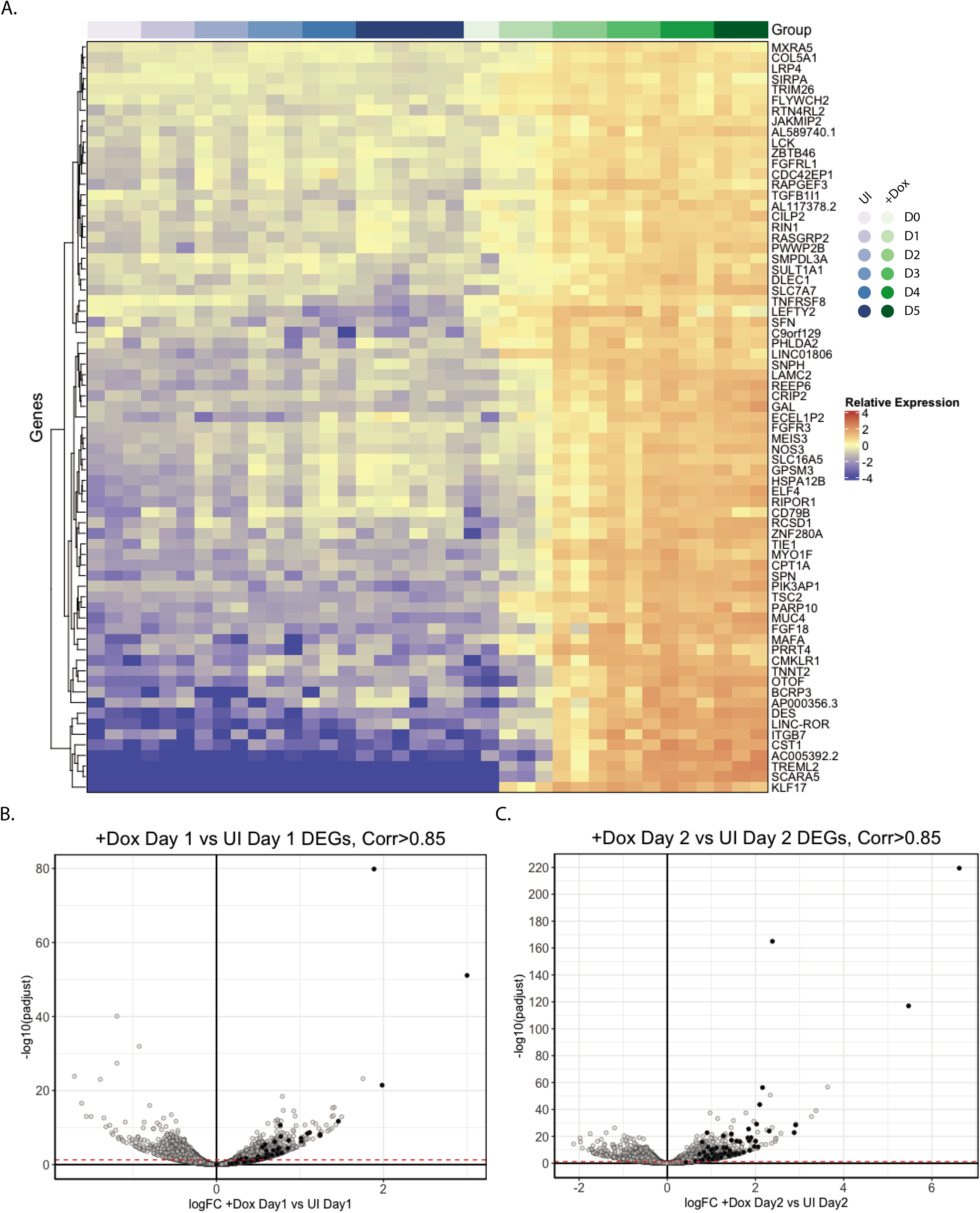
70 genes are strongly correlated with *KLF17* expression over time. **(A)** Heatmap ordered by sample (UI or +Dox) and time point showing all genes that are highly correlated with *KLF17* across time (Pearson correlation coefficient ≥ 0.85). **(B-C)** Volcano plots displaying relative expression of all detected genes in +Dox versus UI H9 KLF17-HA hESCs at (B) day 1 (logFC(+Dox Day1 vs UI Day1)) and (C) day 2 (logFC(+Dox Day1 vs UI Day1)) against the significance of differential expression (-log10(padjust)). The red dotted line notes p_adj_ = 0.05. Individual genes with correlation coefficient to *KLF17* ≥0.85 are displayed as filled circles.

**Figure S3 –.**
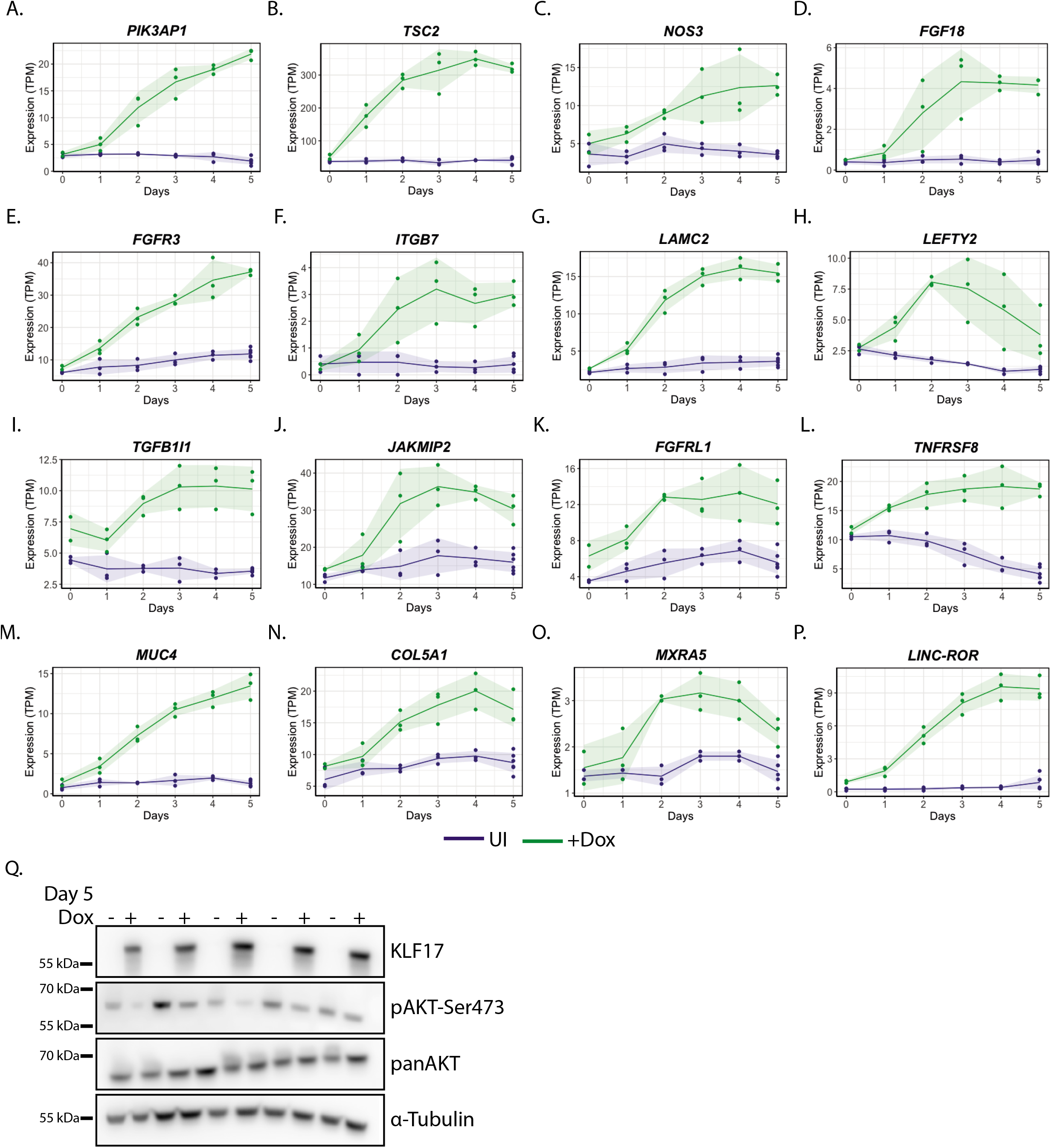
Genes highly correlated with *KLF17* expression include numerous signalling components and cytoskeletal/ECM components. **(A-P)** Normalised expression (TPM) of individual genes of interest across the 5-day time course showing factors involved in (A-G) PI3K-AKT signalling, (H-I) TGFβ signalling, (J-L) other signalling pathways, (F-G,M-O) the cytoskeleton/ICM and (P) the pluripotency-regulating long non-coding RNA *LINC-ROR*. **(Q)** Western blot showing KLF17 induction following 5 days Dox treatment of H9 KLF17-HA and associated expression of PI3K-AKT signalling factors.

**Figure S4 –.**
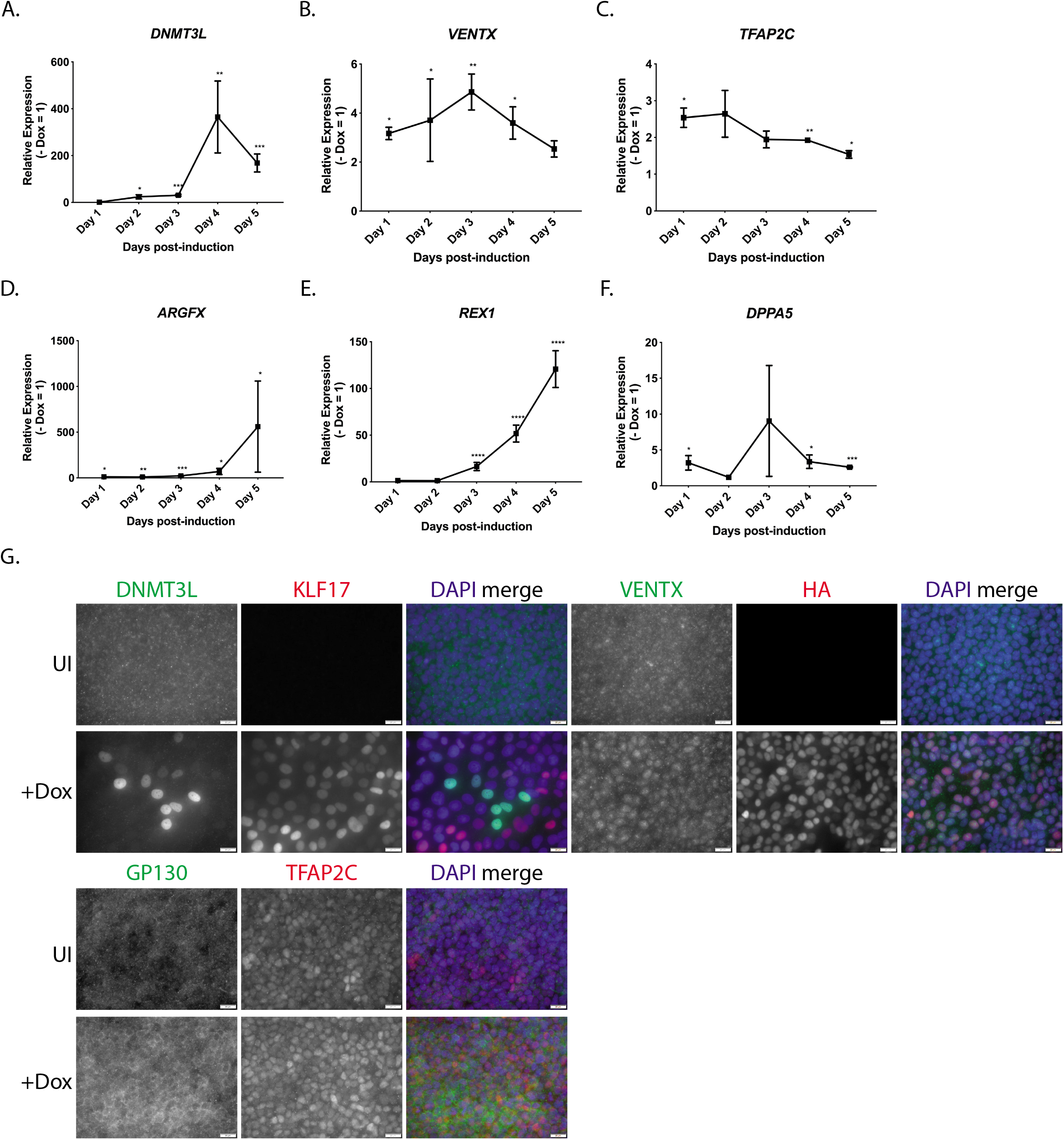
Confirming upregulation of naïve-associated factors following 5 days induction of *KLF17*. **(A-F)** qRT-PCR analysis across the 5-day time course of Dox treatment in H9 KLF17-HA hESCs. Relative expression is displayed as fold change versus uninduced cells and normalised to *GAPDH* as a housekeeping gene using the ΔΔCt method. Dots represent the mean and whiskers the SEM. Welch’s *t* test, **** *p* < 0.001, *** *p* < 0.005, ** *p* < 0.01, * *p* < 0.05. **(G)** Immunofluorescence analysis of H9 KLF17-HA inducible hESCs following 5 days uninduced (UI) or 5 days doxycycline (Dox) induction (+Dox). Scale bars = 20 μm. N ≥ 3.

**Figure S5 –.**
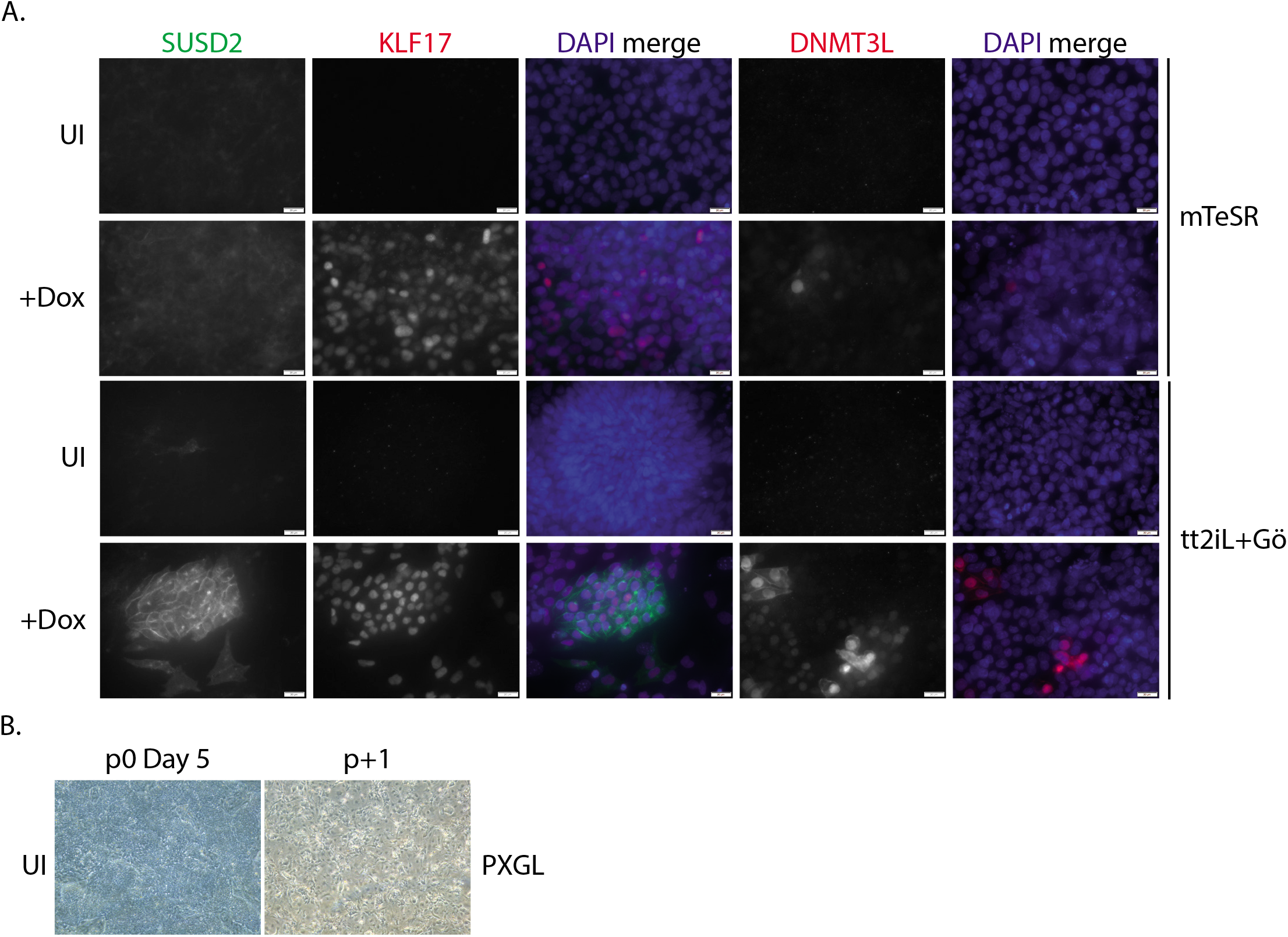
PXGL is uniquely able to support KLF17-driven naïve resetting of H9 KLF17-HA hESCs. **(A)** Immunofluorescence analysis H9 KLF17-HA inducible hESCs following 5 days uninduced (UI) or 5 days doxycycline induction (+Dox) in the indicated media. Cells were cultured on a mouse embryonic fibroblast (MEF) feeder layer and at 5% O_2_. Scale bars = 20 μm. N ≥ 3. **(B)** Unlike induced cells, uninduced control H9 KLF17-HA (UI) were unable to survive in PXGL medium following the first passage.

**Figure S6 –.**
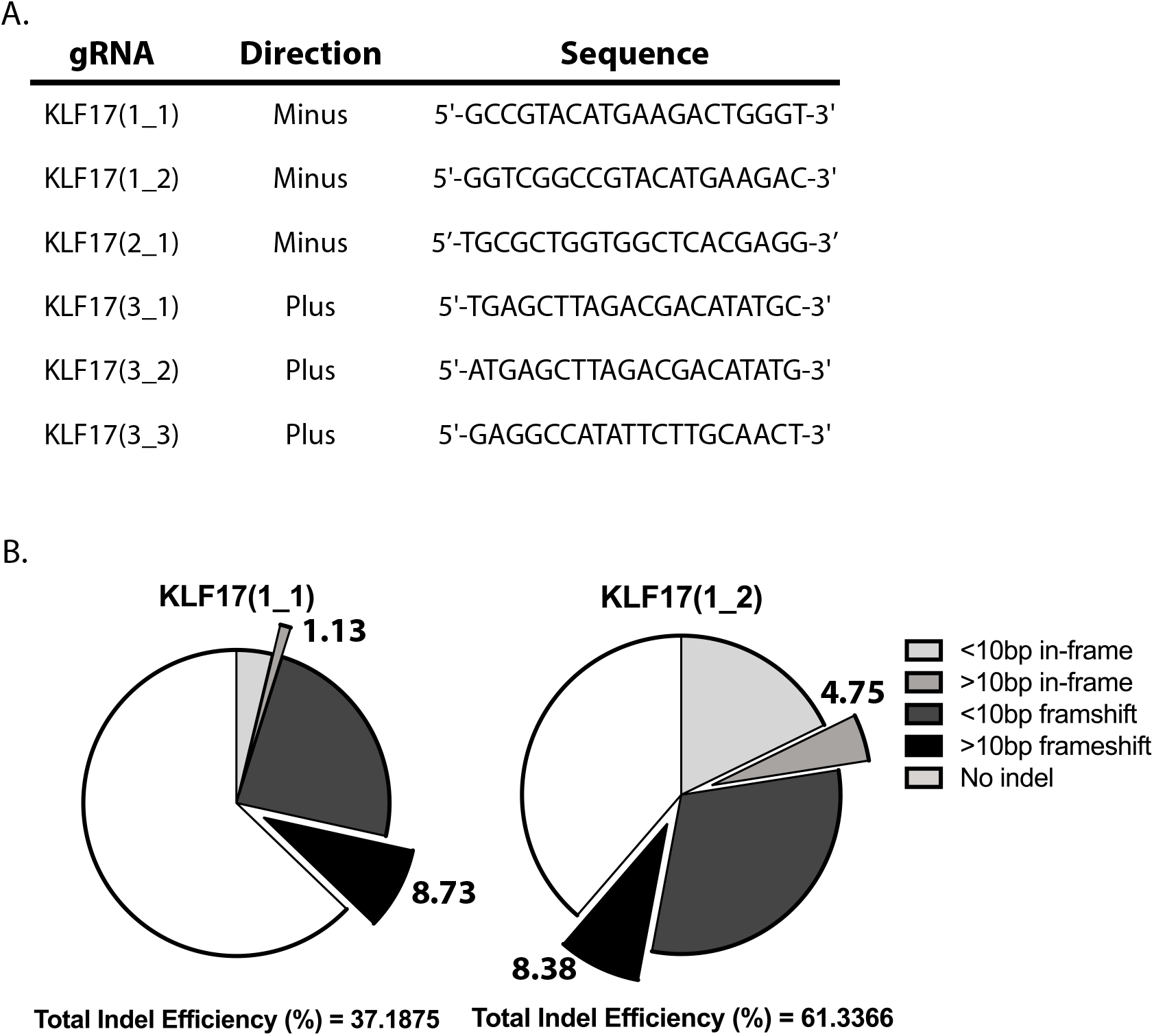
Generating *KLF17*-null hESCs by CRISPR-Cas9. **(A)** A table showing the gRNA sequences tested for mutagenic efficiency. **(B)** Pie charts representing the relative proportions of different outcomes of CRISPR-Cas9 editing of H9 hESCs, based on the sequences detected by MiSeq.

**Figure S7 –.**
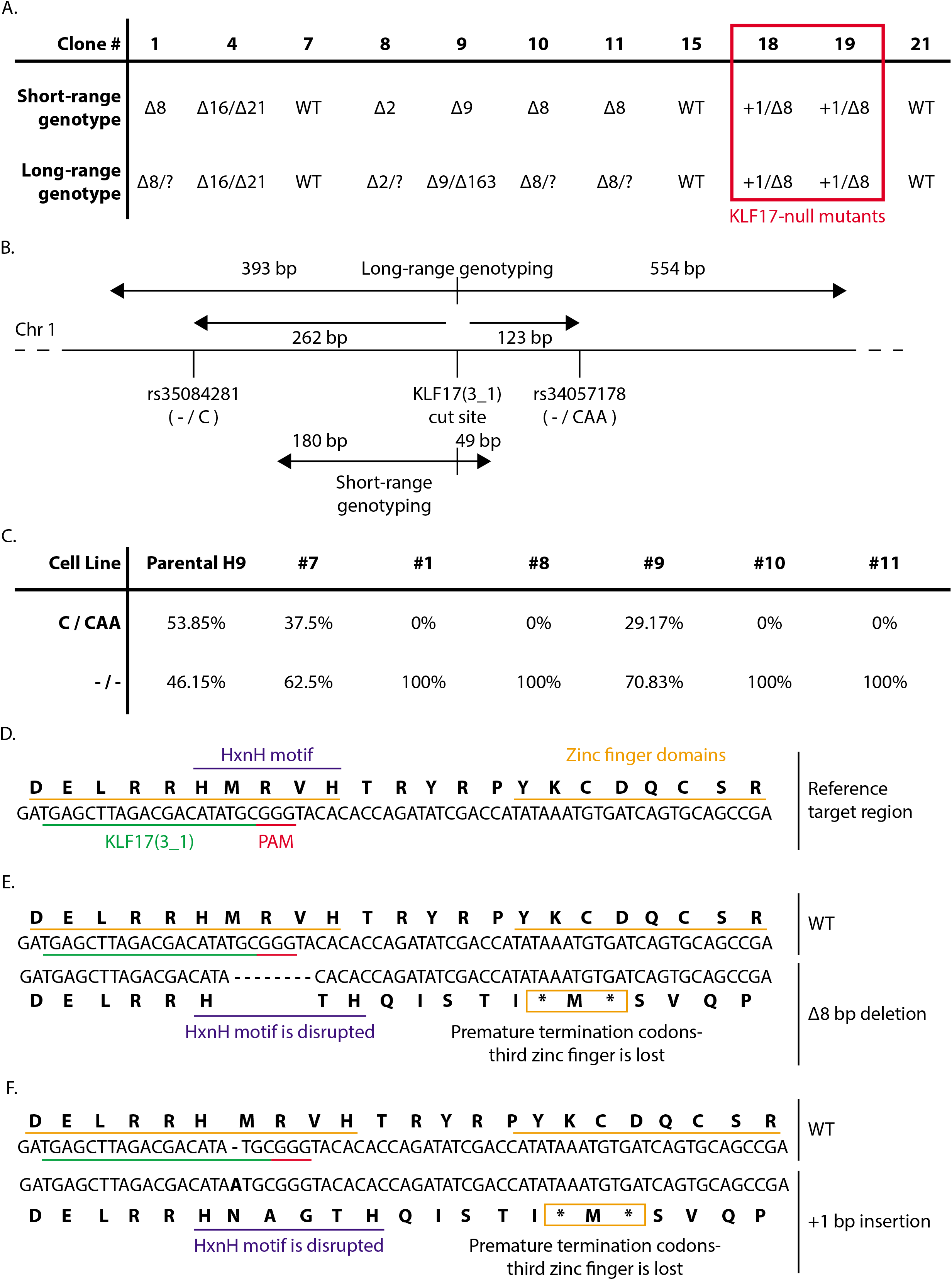
Genotyping of H9 hESCs following targeting with gRNA KLF17(3_1) and clonal expansion. **(A)** A table showing the results of genotyping 11 clones generated following CRISPR-Cas9 targeting. Short-range genotype denotes the results of MiSeq of a ~250 bp region surrounding the KLF17(3_1) cut site. Long-range genotype denotes the results of Sanger sequencing of a ~950 bp region surrounding the KLF17(3_1) cut site. The red rectangle highlights the verified *KLF17^-/-^* H9 hESCs that were carried forward. **(B)** Schematic of the short- and long-range genotyping approach employed on the 11 clones in part (A). **(C)** A table showing the percentage of interpretable reads that showed one of two possible variant-types at the highly polymorphic regions illustrated in (B) - rs35084281 and rs34057178. Parental H9 is the unmodified control cell line, #7 is an internal wild-type control clone generated following nucleofection of KLF17(3_1), #1, #8, #9, 10 and #11 are the KLF17-targeted H9 clones that appeared to have undergone homozygous editing based on short-range genotyping. **(D-F)** Illustration of the sequence context surrounding the KLF17(3_1) cut site in (D) the wild-type reference sequence, (E) the case of an 8 bp deletion and (F) the case of a 1 bp insertion. Important features of the KLF17 sequence are highlighted. DNA sequence is shown in regular font, amino acid sequence is bold above or below the DNA.

**Figure S8 –.**
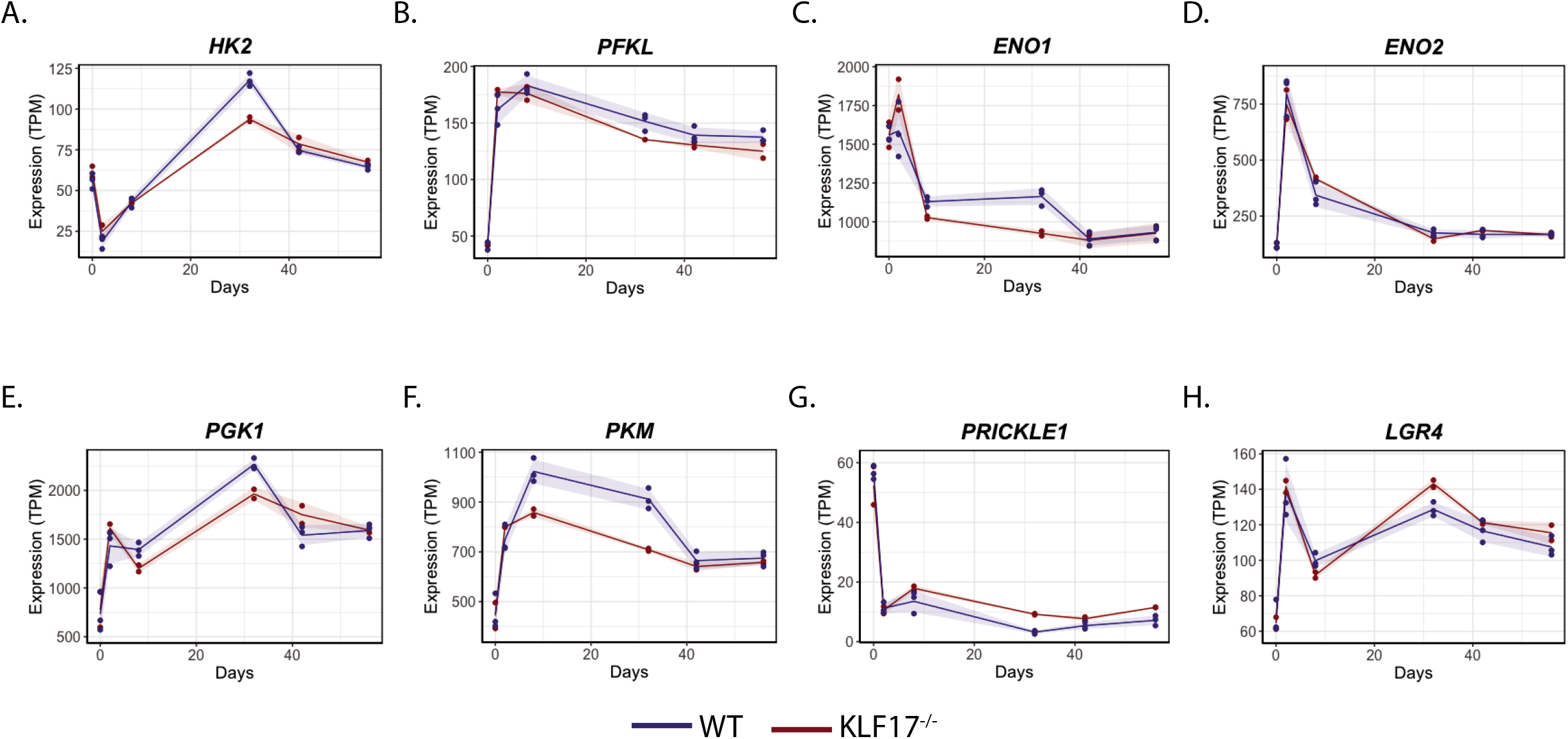
*KLF17*-null naïve hESCs at passage 5 display misregulated expression of core glycolytic enzymes and WNT pathway components. Normalised expression (TPM) of individual genes of interest across the resetting protocol showing (A-F) downregulation of glycolytic enzymes and (G-H) upregulation of WNT signalling factors.

